# Super-resolution histology of paraffin-embedded samples via photonic chip-based microscopy

**DOI:** 10.1101/2023.06.14.544765

**Authors:** Luis E. Villegas-Hernández, Vishesh K. Dubey, Hong Mao, Manohar Pradhan, Jean-Claude Tinguely, Daniel H. Hansen, Sebastián Acuña, Bartłomiej Zapotoczny, Krishna Agarwal, Mona Nystad, Ganesh Acharya, Kristin A. Fenton, Håvard E. Danielsen, Balpreet Singh Ahluwalia

## Abstract

Fluorescence-based super-resolution optical microscopy (SRM) techniques allow the visualization of biological structures beyond the diffraction limit of conventional microscopes. Despite its successful adoption in cell biology, the integration of SRM into the field of histology has been deferred due to several obstacles. These include limited imaging throughput, high cost, and the need for complex sample preparation. Additionally, the refractive index heterogeneity and high labeling density of commonly available formalin-fixed paraffin-embedded (FFPE) tissue samples pose major challenges to applying existing super-resolution microscopy methods. Here, we demonstrate that photonic chip-based microscopy alleviates several of these challenges and opens avenues for super-resolution imaging of FFPE tissue sections. By illuminating samples through a high refractive-index waveguide material, the photonic chip-based platform enables ultra-thin optical sectioning via evanescent field excitation, which reduces signal scattering and enhances both the signal-to-noise ratio and the contrast. Furthermore, the photonic chip provides decoupled illumination and collection light paths, allowing for total internal reflection fluorescence (TIRF) imaging over large and scalable fields of view. By exploiting the spatiotemporal signal emission via MUSICAL, a fluorescence fluctuation-based super-resolution microscopy (FF-SRM) algorithm, we demonstrate the versatility of this novel microscopy method in achieving superior contrast super-resolution images of diverse FFPE tissue sections derived from human colon, prostate, and placenta. The photonic chip is compatible with routine histological workflows and allows multimodal analysis such as correlative light-electron microscopy (CLEM), offering a promising tool for the adoption of super-resolution imaging of FFPE sections in both research and clinical settings.

## Introduction

Histology refers to the study of the structure and organization of the different cell groups within biological organisms by analyzing the microanatomy of tissues. It involves the use of specialized laboratory techniques and instruments to prepare and examine tissue samples in a microscope. In life sciences, histology is of particular importance for several reasons. Firstly, it enables the identification of structural and functional changes during various physiological and pathological processes, including the development of diseases. This information is essential for accurate diagnosis, treatment planning, and monitoring of disease progression. Secondly, histological analyzes permit a better understanding of the mechanisms of disease development and progression, which can lead to the advancement of new and improved treatments. Finally, histology plays a crucial role in many other fields of research, including developmental biology, genetics, and neuroscience, by providing insights into the fundamental structure and organization of the different cells and tissues in plants, animals, and humans.

A standard histological analysis involves several steps including tissue sampling, fixation, sectioning, and labeling, before observation under a microscope. Out of several methods, formalin-fixation paraffin-embedding (FFPE) has become the standard histological processing technique in light microscopy, since it allows an easy, repetitive, reliable, and cost-effective way for preserving, slicing, and archiving tissue samples for decades^1^. Moreover, the FFPE processing method supports several labeling procedures, including light-absorbing dyes such as hematoxylin and eosin (H&E), immunohistochemical markers, and fluorophores. Nowadays, FFPE is estimated to be the most common histological preservation method, with hundreds of millions of samples stored in biobanks around the world^2, 3^. All these features make FFPE a valuable source of biological material for a wide variety of studies ranging from genomics^4, 5^ and proteomics^6^ to aid in diagnosis^1^ and prognosis^7, 8^ of diseases.

High resolution is a desirable feature in histology, as it enables the identification of morphological features relevant both for basic research and clinical purposes. Conventional optical microscopes and slide scanners offer a fast and relatively inexpensive way to observe biological samples, at the cost of limited resolution (∼250 nm), whereas electron microscopes allow for nanoscale resolution at the expense of high operating costs and low throughput. In histopathology, the selection between light and electron microscopy is highly dependent on the level of detail necessary to visualize the structures of interest that are essential to render a clinical diagnosis. While a vast majority of pathologies can be analyzed with conventional diffraction-limited optical microscopy, some other disorders such as minimal change disease^9^, primary ciliary dyskinesia^10^, and amyloidosis^11^, require the high-resolution power of electron microscopy for diagnosis.

Recently, a new set of optical microscopy methods have emerged, allowing sub-diffraction resolution via manipulation of the illumination patterns and/or of the photochemical and photokinetic properties of fluorescently-labeled samples^12^. This new set of techniques referred to as fluorescence-based super-resolution optical microscopy (SRM), or optical nanoscopy, breached the resolution gap between light and electron microscopy, some of them reaching down to sub-20 nm resolution, opening new avenues for the investigation of biological mechanisms under optical instruments. To date, several techniques have emerged under the umbrella of SRM, categorized into four main sub-types, namely, single-molecule localization microscopy (SMLM)^13^, stimulated emission depletion microscopy (STED)^14^, structured illumination microscopy (SIM)^15^, and fluorescence fluctuation-based super-resolution microscopy (FF-SRM)^16^.

The development of optical super-resolution methods has given us a glimpse of its potential impact on histopathological applications, allowing for detailed observations of ultrastructural features on standard FFPE sections^3, 17–20^. However, multiple barriers defer the adoption of super-resolution methods in clinical settings. These include: 1) the limited throughput, in terms of field of view; 2) the susceptibility of the super-resolution methods to refractive index heterogeneity and high labeling density inherently present on FFPE tissues; 3) high operational costs of the existing super-resolution techniques; and 4) the system complexity. For example, SMLM methods support sub-50 nm lateral resolution over relatively large fields of view but typically require thousands of frames for successful image reconstruction^21^. Although SIM approaches require significantly fewer images than SMLM (9 or 15 images for 2D-SIM or 3D-SIM, respectively), commercially available SIM systems are commonly limited to fields of view of approximately 50 µm × 50 µm. Recent approaches such as transmission-SIM^22^ have demonstrated an extended field of view, however, at the cost of a compromised resolution. Moreover, SIM methods are prone to reconstruction artifacts, particularly in the presence of refractive index mismatch^23^, which is a potential issue for heterogeneous samples such as FFPE tissue sections. STED, despite being a reconstruction-free method, is a point-scanning technique with low throughput for scanning the centimeter-scale tissue areas commonly used in histology. Furthermore, the light scattering aberrations experienced by the depletion laser ultimately compromises the lateral resolution achievable with STED in FFPE samples^24^. The FF-SRM comprises a set of statistical methods capable of resolving fine structures out of conventional wide-field image stacks, without the need for special equipment or complex sample preparation. However, FF-SRM methods also face reconstruction challenges due to the high density of fluorescent labels present in tissue samples. Hence, an SRM method capable of addressing these challenges while being compatible with the routinary histological workflow of FFPE samples will prove advantageous for embracing super-resolution histology both in research and clinical settings.

Recently, photonic chip-based optical microscopy has been proposed as a versatile tool for the observation of biological samples, allowing for multimodal high-resolution imaging over large fields of view^21, 25–27^. The method consists of a photonic integrated circuit (in short, a photonic chip), that holds the sample while providing it with the necessary illumination for fluorescence microscopy. The photonic chip employed in this work is composed of a silicon substrate layer of silicon (Si), an intermediate layer of silicon dioxide (SiO_2_), and a top optical waveguide core layer of silicon nitride (Si_3_N_4_) that transmits light in its visible spectrum^26^ (Figure 1a). For microscopy analysis, the biological sample is placed in the imaging area (Figure 1b) and further prepared for fluorescence labeling. Thereafter, an excitation beam source is coupled to a chosen waveguide facet, allowing for confined light propagation along the core material via total internal reflection. Upon coupling, an evanescent field with a penetration depth of <50 nm for the chosen photonic-chip forms at the waveguide-sample interface, exciting the fluorescent molecules in its reach. The waveguide geometries support multi-mode interference (MMI) patterns that provide a semi-stochastic, non-uniform illumination to the sample^25, 28, 29^. To achieve isotropic illumination, the MMI patterns are modulated by translating the coupling objective along the input facet of the chip, while individual frames are collected (see Supplementary Video V1 and Supplementary Information S1). For multicolor imaging, the process is repeated using the specific excitation wavelength for each fluorescent marker. Finally, the acquired image stacks are computationally averaged, pseudo-colored, and merged to obtain a chip-based total internal reflection fluorescence (chip-TIRF) image. Contrary to the conventional TIRF that uses a high numerical aperture (NA) and high magnification lenses (usually 60X to 100X, and 1.49 NA) to generate the evanescent fields, the evanescent field here is generated by a photonic chip. Thus, by using conventional microscope objectives of diverse magnifications (see Supplementary Information S2), the photonic chip enables high-contrast TIRF images over diverse fields of view (Figure 1c).

**Figure 1.**
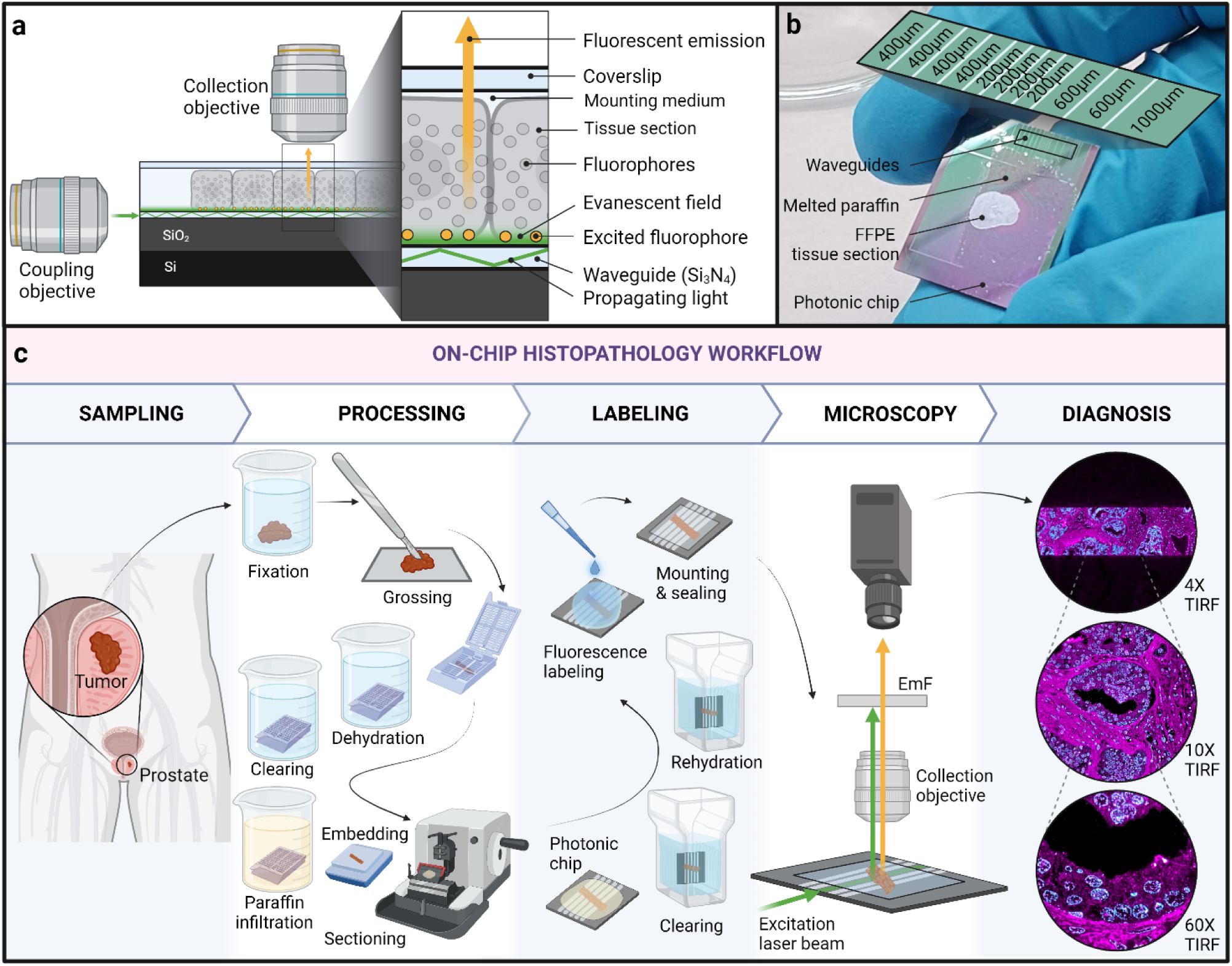
Photonic chip-based microscopy for FFPE samples. **a)** Working principle: upon coupling, the excitation laser beam propagates along the optical waveguide due to the phenomenon of total internal reflection. An evanescent field of <50 nm forms at the waveguide surface, exciting the fluorophores on its reach. The fluorescent emission is collected by a conventional microscope objective lens. **b)** View of a photonic chip containing an FFPE tissue sample after oven incubation at 60 °C for paraffin melting. The zoomed-in area illustrates diverse waveguide widths available on the photonic chip (200 µm, 400 µm, 600 µm, and 1000 µm). **c)** Photonic chip-based microscopy is compatible with standard histological workflows: following extraction from the diseased organ, the tissue sample is further processed via FFPE steps including fixation in formalin, grossing, dehydration, clearing, paraffin infiltration, and embedding. Thereafter, the tissue block is sliced on a microtome into a thin section (2 µm - 4 µm). Subsequently, the tissue section is floated in a water bath and then scooped onto a photonic chip for further paraffin melting, clearing, rehydration, and fluorescence labeling. After mounting and sealing, the chip is placed on a standard upright microscope equipped with a side excitation laser beam. Upon coupling onto a chosen waveguide, the fluorescent signal is collected by a conventional microscope objective while the excitation light is further blocked using an emission filter (EmF). By transitioning from low to high magnification collection objectives, photonic chip-based microscopy offers high-contrast and super-resolution visualizations of FFPE tissue samples over scalable fields of view, facilitating the histological interpretation and subsequent diagnosis of diseases.

In a previous study^30^, photonic chip-based microscopy was shown as a multimodal platform for super-resolution imaging of tissue sections prepared using Tokuyasu cryopreservation method^31^. Particularly, the implementation of photonic chip-based FF-SRM via multiple signal classification algorithm^32^ (MUSICAL) revealed promising potential as a fast super-resolution imaging method for tissue sections. However, previous efforts were somehow limited in throughput and did not cover standard FFPE tissue sections.

In this study, we propose photonic chip-based optical microscopy as a high-contrast super-resolution tool for the observation of FFPE samples over large fields of view. By using FFPE-processed samples of human organs with diverse clinical conditions, namely, colorectal cancer, prostate cancer, and healthy placenta, we demonstrate full compatibility of the photonic chip with conventional histological workflows, including the incubation steps for deparaffinization, re-hydration, and labeling necessaries for fluorescence imaging (Figure 1c). Furthermore, by exploiting the evanescent field excitation offered by the chip, we show the superior contrast and super-resolution capabilities of the chip-based FF-SRM and highlight the interplay between the contrast and the resolution offered by the photonic chip across scalable magnifications. We further demonstrate the multimodality supported by the photonic chip for high-resolution correlative light and electron microscopy. To the best of our knowledge, this is the first study of FFPE sections on a photonic chip-based microscope and the first report of FF-SRM on FFPE sections. Photonic chip-based optical microscopy has the potential to assist in high-contrast and high-throughput super-resolution fluorescence imaging of paraffin-embedded samples, paving the road for super-resolution histology both in research and clinical settings.

### Contrast enhancement approaches for histology

Contrast is an essential parameter for the visual interpretation of microscopy images. Broadly speaking, contrast is the difference between the sample signal and its surroundings, either in terms of intensity or color. In histology, a high contrast allows for clear visualization of features of interest located at the microscope objective’s focal plane. However, most tissue samples are translucent, which makes it difficult to observe them under a light microscope. There are diverse mechanisms to add contrast to histological samples. These are divided into two categories, namely, label-free and histochemical methods. The first group exploits physical phenomena such as light interference in phase-contrast computed tomography^33^ and differential interference contrast^34^; or light scattering in Raman spectroscopy^35^, darkfield and optical coherence tomography^36^; or nonlinear optics in second harmonic generation^37^ to retrieve high contrast information from the unlabeled samples. The latter group employs either light-absorbing or light-emitting compounds to chemically stain specific areas of the tissues to improve their visualization under the microscope.

Traditionally, the histology field has relied on light-absorbing dyes for the pigmentation of tissue samples. The most popular staining, hematoxylin and eosin (H&E), enables the distinction between cell nuclei, extracellular matrix, and cytoplasmic content within the tissues^38^. In addition, the immunohistochemical techniques improve the labeling specificity as compared to H&E, supporting low contrast identification of protein expression within the tissues. In combination with immunogenic approaches, the fluorescent markers also enable high labeling specificity, which makes them suitable for diverse analyses such as multiplexing^39^, quantitative tissue cytometry^40^, and image segmentation^41^, among others. Nonetheless, the contrast ratio of fluorescent signals is greatly affected by the out-of-focus blur arising from emitting fluorophores located in the foreground and/or the background of the focal plane. Therefore, removing the off-focus information is essential for an accurate interpretation of the fluorescent data. This can be achieved in three ways:

a. **By mechanically slicing the samples into thin sections.** Typically, the thinnest section thickness for FFPE samples is approximately 2 µm – 4 µm, which is insufficient for reducing the out-of-focus signal of fluorescently-labeled tissue samples. To obtain even thinner tissue sections (down to 70 nm thickness), alternative histological approaches, such as ultrathin sectioning of resin-embedded or cryopreserved samples, have been proposed.
b. **By optical sectioning.** By manipulating the excitation and/or emission light paths to selectively illuminate and collect information from specific planes of the sample, a method called optical sectioning, it is possible to minimize the out-of-focus information. Examples of these are confocal microscopy^42^, multi-photon excitation microscopy^43^, total internal reflection fluorescence (TIRF) microscopy^44^, and light-sheet microscopy^45^.
c. **By image post-processing.** A new set of computational methods have emerged as an appealing alternative to enhance contrast by artificially removing the blurring effects caused by the out-of-focus light and the optical aberrations of the microscope system. These methods, based on mathematical algorithms such as deconvolution^46, 47^, structured illumination^15^, computational clearing^48^, and machine learning^49^, greatly improve the contrast and assist in restoring the latent high-resolution signal out of the observed low-resolution fluorescence data.

### Proposing TIRF for high-throughput FF-SRM histology

Contrast plays a key role in fluorescence super-resolution optical microscopy, as it defines the extent to which fine details can be observed. From a physics perspective, the optical transfer function of the imaging system acts as a low-pass filter that favors the visualization of coarse features, at the expense of a low-contrast view of the finer structures in the sample. The characteristic triangular shape of the optical transfer function dictates the contrast achievable at diverse spatial frequencies present in the sample^50^. Thus, low spatial frequencies (larger sample features) benefit from high contrast, while high spatial frequencies (smaller sample features) experience low contrast. Moreover, the recorded grayscale intensity of high spatial frequencies often matches the base level of diverse high-frequency system disturbances such as Poisson noise, electronic noise, and dark noise, which makes it even more difficult to differentiate small sample features from noise^51^.

Consequently, contrast enhancement strategies are frequently implemented to improve the performance of the super-resolution methods. Among these, TIRF is an attractive route for achieving high contrast fluorescence on biological specimens, as it limits the axial illumination to a thin layer that provides optical sectioning at the interface between substrate and sample^52^. In recent studies, TIRF illumination was used in combination with single-molecule localization to achieve sub-diffraction views of FFPE samples^3, 20^. However, the fields of view obtained in these studies (roughly, 50 µm × 50 µm) are too small for routine histological analysis. Although a novel prism-based SMLM approach recently showed 40 nm resolution over a half-millimeter field of view^53^, like conventional SMLM approaches, the method required a large number of frames (30,000) to reconstruct a single super-resolved image. These two limitations make SMLM unattractive for high-throughput imaging of FFPE samples. Albeit not achieving as good resolution as SMLM, FF-SRM requires significantly less number of frames (usually hundreds but as low as 30 frames have been demonstrated^54^). Thus, the combination of FF-SRM with the excellent optical sectioning provided by TIRF illumination is an attractive route for high-speed imaging of FFPE tissue sections.

## Results

### Enabling FF-SRM histology via chip-TIRF modulation

In this part of the study, we used a human placental tissue section to demonstrate the superior performance of the photonic chip-based TIRF (chip-TIRF) over epifluorescence (in short, EPI) illumination for achieving FF-SRM on FFPE samples. Here, a chorionic villi sample was taken from the fetal side of a human placenta, dissected, and further embedded in paraffin following a standard FFPE method. Thereafter, the tissue block was sectioned into a 3 µm slice and placed on a photonic chip for further histological processing (see detailed preparation protocols in the Materials and Methods section). After fluorescence labeling, two image stacks (500 frames each) were collected over the same field of view using a 60X/1.2NA water immersion objective under EPI and chip-TIRF, respectively.

The upper segment in Figure 2a illustrates a single EPI image of an FFPE human placental tissue section. Using EPI modality, the microscope objective is utilized both for the excitation and the collection of the fluorescent signal. In the acquisition process, the excitation light beam interacts with the whole sample volume, illuminating all the fluorophores along its path, as shown in Figure 2c. Consequently, the microscope objective captures the fluorescent emission from the illuminated volume, causing out-of-focus blur. In photonic chip-based microscopy (Figure 2d), on the contrary, only the bottom part of the sample is illuminated, enabling ultrathin optical sectioning <50 nm that improves the contrast of the fluorescence microscopy image (upper segment in Figure 2b) as compared to EPI.

**Figure 2.**
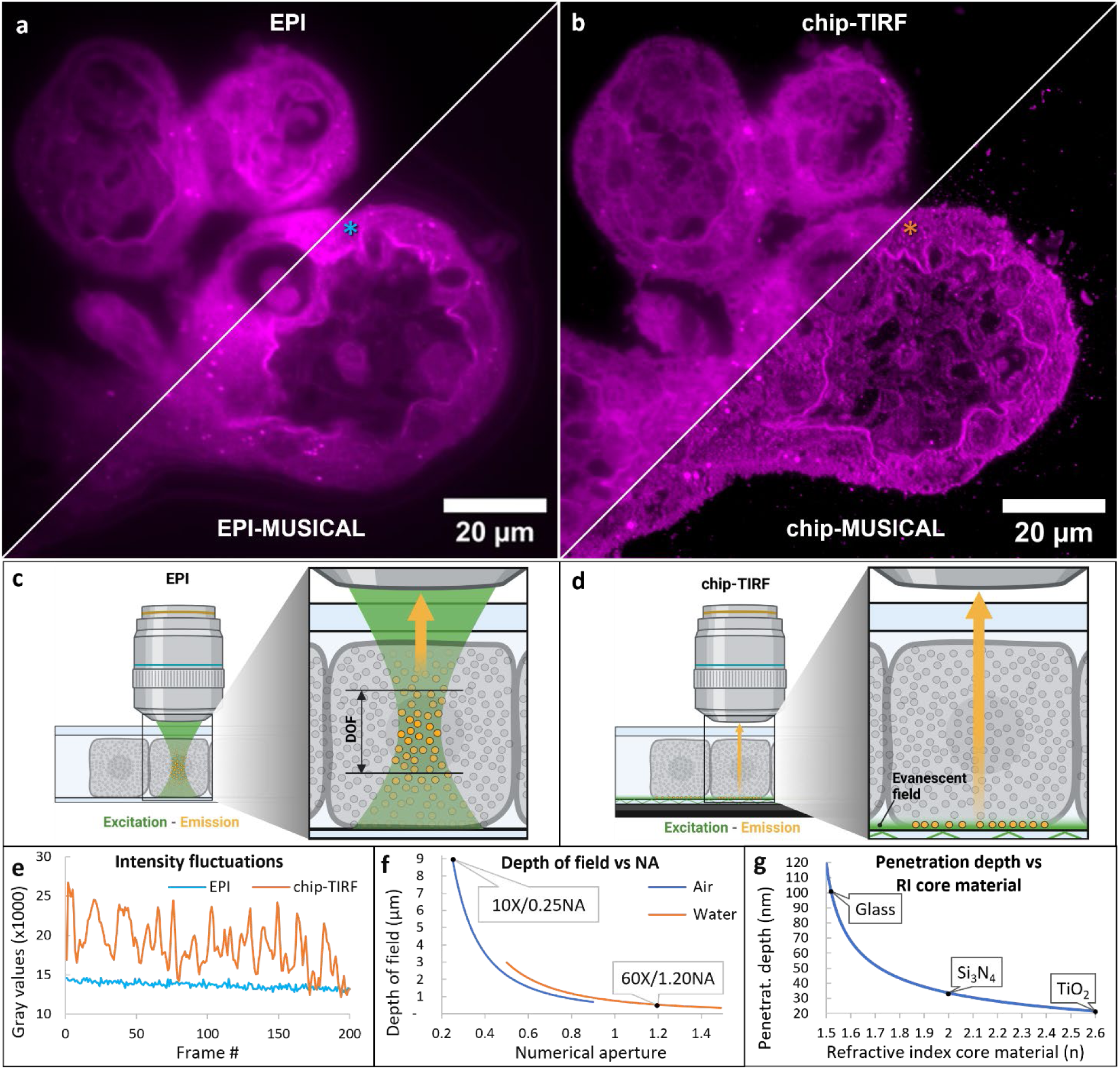
FF-SRM comparison on a human placental FFPE tissue section via EPI and chip-TIRF illumination. **a)** The upper segment illustrates an EPI fluorescence image of the tissue sample. The lower segment shows the EPI-based MUSICAL reconstruction. The blue asterisk illustrates a single pixel chosen for plotting the intensity fluctuations in e). **b)** The upper segment illustrates a diffraction-limited chip-TIRF image. The lower segment shows the chip-based super-resolved image via MUSICAL. The orange asterisk illustrates a single pixel chosen for plotting the intensity fluctuations in e). **c)** Schematic representation of an EPI fluorescence image acquisition. In EPI modality, the microscope objective lens is used both for the illumination and collection of the fluorescent signal. The fluorescent signal is collected from a sample volume corresponding to the depth of field (DOF) of the objective that, for densely-labeled samples such as FFPE sections, results in a highly-averaged signal with low frame-to-frame variance. **d)** In chip-TIRF, the sample is illuminated via evanescent field excitation, allowing for ultrathin optical sectioning that minimizes the signal averaging issues while providing superior spatial and temporal variance that contribute to optimal FF-SRM reconstructions via MUSICAL. **e)** Intensity fluctuations of a single pixel over 200 frames of EPI and chip-TIRF image stacks. Chip-TIRF provides higher temporal variance compared to EPI. See Supplementary Video V2 for details. **f)** Theoretical simulation of DOF vs numerical aperture of air and water-immersion objective lenses. While in EPI the DOF leads to signal averaging, in photonic chip-based microscopy the illumination and the collection light paths are decoupled, which allows for FF-SRM irrespective of the DOF of the collection objective. See Supplementary Information S3 for details. **g)** Theoretical simulation of penetration depth vs refractive index. The penetration depth of TIRF systems is highly dependent on the refractive index contrast between the sample media (n ≈ 1.4) and the core material used for TIRF. The waveguide material (Si_3_N_4_, n ≈ 2) used in this work enables superior optical sectioning capabilities (<50 nm), compared to existing glass-based TIRF approaches employing materials with lower refractive index (borosilicate, n ≈ 1.52). Alternative waveguide materials such as titanium dioxide (TiO_2_, n ≈ 2.6) can potentially improve the optical sectioning to around 20 nm. See Supplementary Information S4 for details.

To further compare the performance of EPI and chip-TIRF under FF-SRM, we chose the multiple signal classification algorithm^32^ (MUSICAL). Through singular value decomposition, this method achieves super-resolution by discriminating between signal and noise spaces across a given diffraction-limited fluorescence image stack. In addition, MUSICAL enables super-resolution imaging even in situations of low excitation intensities, fast acquisition, and relatively small datasets. Here, we post-processed the previously collected EPI and chip-TIRF stacks with MUSICAL. The results are illustrated in the lower segments of Figure 2a and Figure 2b, respectively. Although the EPI-MUSICAL reconstruction resulted in neither noticeable resolution nor contrast enhancement compared to the diffraction-limited EPI fluorescence image, chip-MUSICAL improved both the resolution and contrast when compared to the diffraction-limited chip-TIRF image. This improvement enabled a clearer visualization of the placental tissue structure. The discrepancy in MUSICAL performance observed here can be attributed to the difference in fluorescence intensity fluctuations, both spatially and temporally, between the EPI and chip-TIRF imaging modalities. Specifically, the FF-SRM algorithms require high signal variance among adjacent fluorophores to successfully perform statistical analysis of the fluorescent data within an image stack. Typically, the temporal variance is derived from the gray value fluctuations occurring within consecutive frames, while the spatial variance is achieved through sparse labeling and/or illumination of the sample. As further elaborated herein, unlike EPI, the combination of MMI pattern modulation and ultrathin optical sectioning supported by the photonic chip enables the necessary fluorescence intensity fluctuations for optimal FF-SRM reconstruction of FFPE tissue sections.

In EPI, the temporal fluctuations are determined by the stochastic emission of the fluorophores. Therefore, short acquisition rates are required to minimize the averaging effects of the camera exposure time. In photonic chip-based microscopy, on the contrary, the temporal fluctuations are achieved by modulating the MMI illumination patterns on a frame-to-frame basis, which allows for longer camera exposure times. Here, this aspect is illustrated by plotting the gray values of an arbitrary pixel over the collected image stacks (blue and orange asterisks in Figure 2a and Figure 2b, respectively). Despite the longer acquisition time of the chip-TIRF modality (50 ms per frame) as compared to the short camera exposure time of the EPI modality (10 ms per frame), the photonic chip-based method enabled over 30 % frame-to-frame variability (orange line in Figure 2e), whereas EPI supported, at most, 7 % variability at the same sample location (blue line in Figure 2e). Arguably, the low variance of EPI compromised the statistical analysis of the chosen FF-SRM algorithm. The chip-TIRF stack, in turn, resulted in optimal data for FF-SRM reconstruction, as reported in previous observations^25, 28–30^. Supplementary Video V2 provides a detailed view of the fluorescence fluctuations obtained in EPI and chip-TIRF modalities.

Another important parameter for successful FF-SRM reconstruction is spatial sparsity. Although FF-SRM algorithms are designed for multi-emitter datasets, these methods perform best for signals where the information is sparsely distributed in the spatial domain. The signal sparsity can be achieved by means of sparse sample labeling and/or by modulation of the illumination source. In the case of the relatively thick and densely labeled FFPE sections, spatial sparsity can be achieved by combining random illumination and thin optical sectioning. While the first can be implemented in EPI via speckle illumination^55^, the optical sectioning capabilities of this imaging modality are limited by the depth of field (DOF) of the microscope objective (Figure 2c and Figure 2f), resulting in an averaged signal with low spatial sparsity. In chip-TIRF modality, on the contrary, the sample is illuminated by an evanescent field that restricts the fluorescence emission to a thin sample layer (Figure 2d and Figure 2g) and semi-stochastic MMI patters that allow both for ultrathin optical sectioning and for random illumination. In the placental section, for example, despite the relatively short DOF (approx. 520 nm) used for EPI, the abundance of fluorophores within the excited sample volume resulted in a dense signal with low spatial sparsity that complicated the EPI-MUSICAL reconstruction. In chip-based microscopy, however, the combined optical sectioning and random illumination supported by this technique successfully enabled chip-MUSICAL reconstruction. Supplementary Information S3 provides detailed information about the DOF of the diverse microscope objectives used in this study.

The high refractive index of the waveguide core material (*n* ≈ 2, for Si_3_N_4_) provides additional advantages for histology. Firstly, it allows for high spatial frequencies in the illumination that prove advantageous for improved lateral resolution of other SRM methods such as SMLM^25^ and SIM^56^. Secondly, it limits the penetration depth of the evanescent field below 50 nm for our chosen photonic chip design (Figure 2g), allowing superior optical sectioning compared to glass-based TIRF approaches employing lower refractive index materials such as borosilicate glass (*n* ≈ 1.52). Furthermore, alternative waveguide materials with even higher refractive index such as titanium dioxide (TiO2, n ≈2.6) could potentially improve the optical sectioning to around 20 nm (see Supplementary Information S4 for details). In the next section, we further investigate how chip-TIRF assists in the histological assessment of FFPE tissue sections.

### Evanescent field excitation for high-contrast histology

In this part of the study, we used an FFPE-preserved colorectal cancer sample to demonstrate two features of the photonic chip-based microscopy: a) its compatibility with standard histochemical processing techniques, and b) the superior contrast capabilities offered by this novel technique as compared to other fluorescence-based microscopy methods. To achieve this, we first explored the transition from conventional hematoxylin and eosin (H&E) staining to fluorescent labeling and, subsequently, we focused on the contrast performance of the photonic chip-based microscopy.

Here, an FFPE colorectal sample was sectioned into two consecutive slices of approx. 1 cm × 1 cm × 3 µm (width, height, thickness) and subsequently floated in a water bath (see detailed preparation protocols in the Materials and Methods section). For conventional histology, one slice was mounted on a glass slide and histochemically prepared following H&E staining procedures. For fluorescence imaging, the remaining slice was mounted on a photonic chip and further stained with fluorescent markers against membranes and nuclei. The photonic chip withstood all the harsh steps of the histochemical processing (see Supplementary Information S5), including water immersion for sample scooping (see Supplementary Video V3), 60 °C oven incubation for paraffin melting, xylene baths for paraffin clearing, sample rehydration through descendent ethanol series, and fluorescent labeling. In addition, we successfully tested antigen retrieval steps on a different paraffin-embedded sample (see Supplementary Information S6) by microwave boiling the tissue section on the chip before immunolabeling.

The H&E-stained sample was imaged on a whole slide scanner device (Virtual Slide System V120, Olympus) using a 20X/0.85NA objective lens. The images were automatically stitched to obtain a complete view of the sample, as shown in Figure 3a. The hematoxylin, shown in purplish hues/colors, allowed the identification of nuclei whereas the eosin, in shades of pink, facilitated the observation of the cytoplasmic content and extracellular matrix. Here, the H&E staining enabled the identification of four major microanatomical areas in the colorectal sample (black-dotted lines in Figure 3a), namely, smooth muscle (Sm), necrotic tissue (Ne), benign colonic epithelium (Be), and adenocarcinoma (Ad). The identification of the diseased sections was carried out by the pathologist. While a zoom- in view of the adenocarcinoma region is sufficient to reveal the cancer cells forming glands (inset in Figure 3a), a higher resolution and contrast could further aid the visualization of the diagnostic features (see Supplementary Information S7).

**Figure 3.**
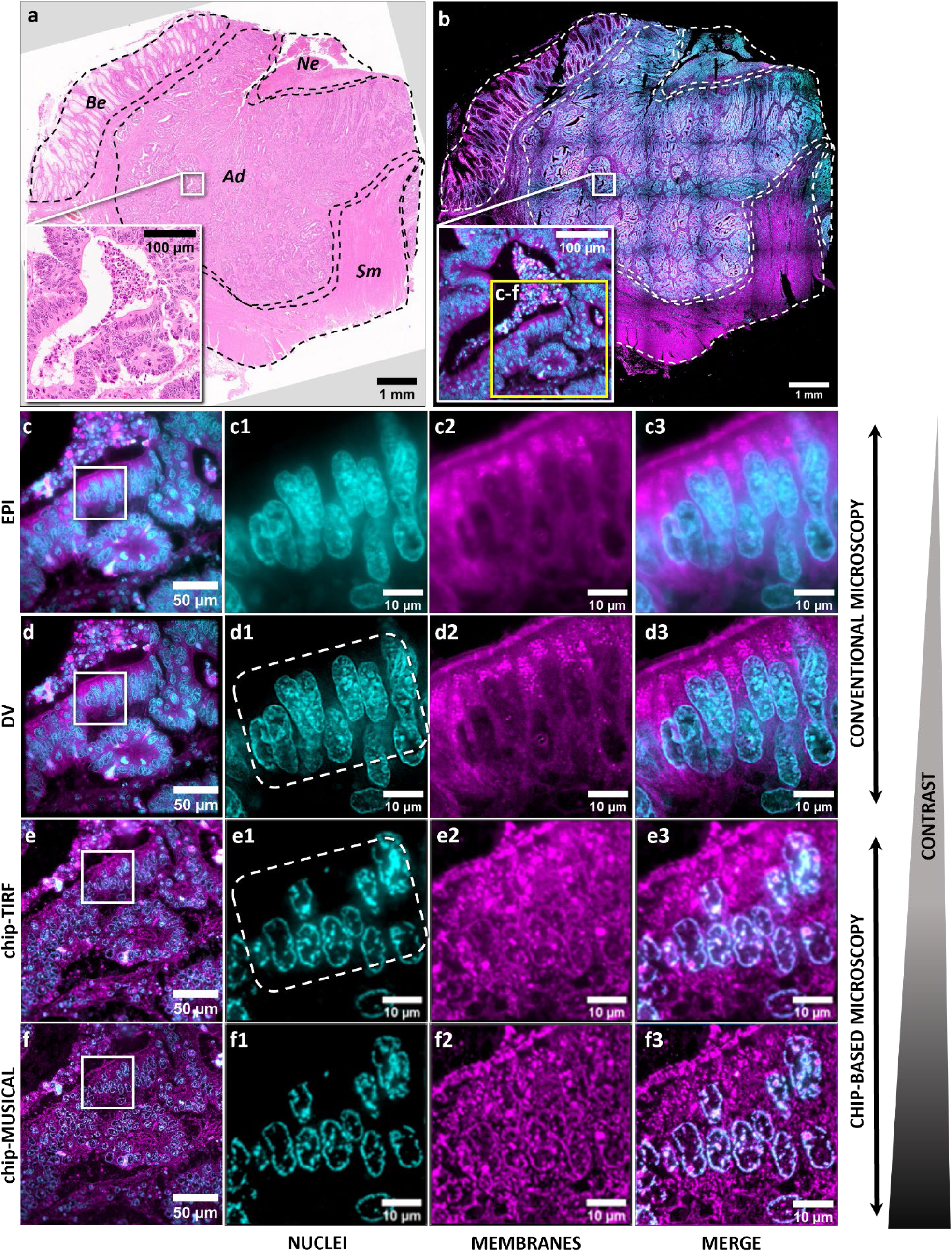
High-contrast histology via photonic chip-based total internal reflection fluorescence microscopy. **a)** Bright-field image of a human FFPE colorectal cancer tissue section stained with hematoxylin and eosin. Back-dotted lines denote four major regions within the sample, namely, smooth muscle (Sm), necrotic tissue (Ne), benign colonic epithelium (Be), and adenocarcinoma (Ad). A close-up view of the Ad area allows for the identification of gland-forming tumor cells, with nuclei in purple and cytoplasm content in pink. **b)** Multicolor EPI fluorescence image of a consecutive slice, labeled with fluorescent markers against nuclei, and membranes, and pseudo-colored in cyan and magenta, respectively. The grid-like artifact is a consequence of the non-uniform Gaussian profile characteristic of EPI illumination. The fluorescent labeling not only improves the contrast between nuclei and membranes, as shown in the close-up box, but also enables a clearer distinction among the four regions of the sample compared to H&E. For example, between the Ad, the Ne, and the Sm areas. **c-f)** 60X view of the yellow box in b) under various fluorescence-based microscopy methods. **c)** EPI fluorescence microscopy image of a region of interest within the Ad. Close-up views **c1** and **c2** illustrate the nuclei and membrane channels, respectively. The merged view in **c3** exhibits a characteristic blur inherent to EPI illumination. **d)** Deconvolution microscopy image of the same region of interest in Ad. Close-up views **d1**, **d2**, and **d3** show a significant contrast improvement compared to their corresponding EPI images. However, a detailed view inside the white-dotted line in d1 reveals the superposition of the nuclear signal that hampers the quantification of these structures. **e)** Photonic chip-based microscopy (chip-TIRF) offers ultrathin optical sectioning of the colorectal sample, enabling contrast improvement both in the lateral and axial domains. Particularly, individual nuclei can be easily identified inside the white-dotted line in e1, enabling the quantification of such structures. **f)** Statistical post-processing of the photonic chip-based microscopy raw data via fluorescence fluctuation-based SRM algorithms such as MUSICAL (chip-MUSICAL) further improves the contrast and resolution of the chip-TIRF images, enabling a detailed view of the sample. Chip-based microscopy acquired with a 60X/1.2NA provides superior contrast as compared to conventional EPI fluorescence-based methods such as EPI and DV acquired with a 60X/1.42NA.

Similarly, the on-chip colorectal sample was imaged on a commercial EPI fluorescence microscope (DeltaVision Elite Deconvolution Microscope, GE Healthcare) using a 10X/0.4NA objective lens. For each field of view, two consecutive images were taken using different fluorescent channels, namely, far-red channel for membranes, and green channel for nuclei (see details in the Materials and Methods section). Upon pseudocoloring and merging, the 10X EPI fluorescence images were stitched in an 8 × 8 tile mosaic to enable full visualization of the sample. The resulting image (Figure 3b) offers a full visualization of the fluorescently-labeled colorectal sample, with membranes shown in magenta and nuclei shown in cyan. Here, the grid-like artifact on the mosaic image is largely due to the non-uniform Gaussian profile characteristic of EPI illumination, which is further illustrated in Figure 4. Notably, the fluorescent labeling improves the contrast between the labeled structures, allowing for a much clearer distinction between the diverse regions of interest in the sample (white-dotted lines in Figure 3b), as compared to the H&E slide (Figure 3a). For example, in Figure 3b, the smooth muscle (Sm) region shows a dominant magenta color as compared to the neighboring adenocarcinoma (Ad) part, indicating a different distribution of nuclear content between these two regions. Similarly, it is possible to observe a higher density of nuclei (in cyan color) on the necrotic region (Ne) as compared to the other regions in the sample. Arguably, these differences are much more difficult to spot on the low-contrast view of the H&E sample, as illustrated in Figure 3a.

**Figure 4.**
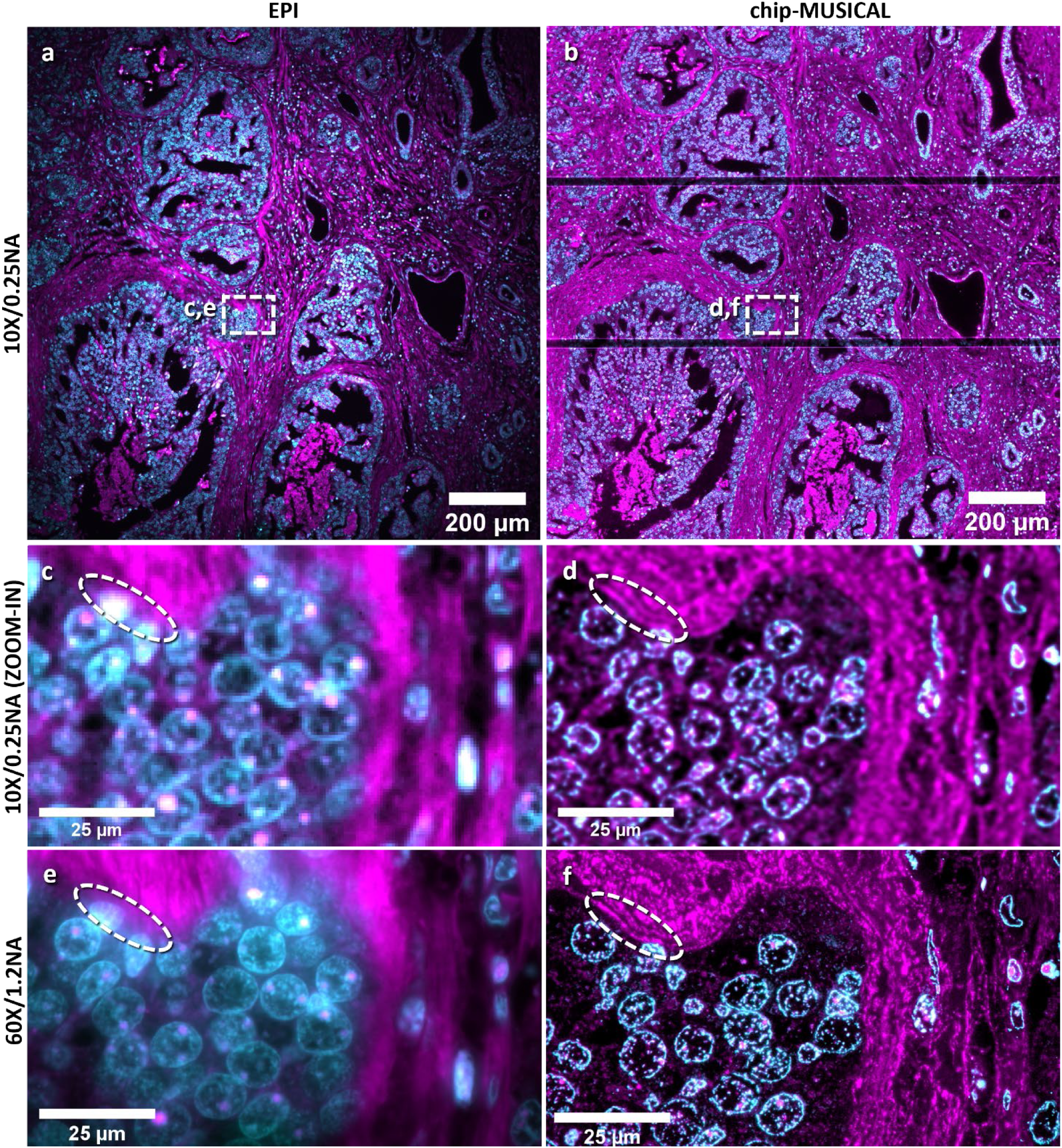
Super-resolution chip-based histology over scalable fields of view. **a)** EPI fluorescence image of an FFPE prostate cancer sample acquired in low magnification with a 10X/0.25 objective lens. Nuclei are shown in cyan and membranes in magenta. The vignetting pattern around the borders of the image is due to the non-uniform Gaussian profile characteristic of EPI illumination. The white-dotted box represents the region of interest further discussed in c) and e). **b)** Same sample area as in a) imaged with chip-based TIRF illumination using a 10X/0.25 objective lens and further analyzed with MUSICAL. The dark horizontal lines denote the location of 25 µm-wide spacing gaps between adjacent waveguides. The white-dotted box represents the region of interest further discussed in d) and f). **c)** Zoom-in view of the white-dotted box segment in a). **d)** Zoom-in view of the white-dotted box segment in b). The white-dotted oval in the 10X chip-MUSICAL image denotes the location of membrane structures that are otherwise not distinguishable in the same region of the 10X EPI fluorescence image. **e)** EPI fluorescence image of the white-dotted box segment in a) acquired in higher magnification using a 60X/1.2NA objective lens. Despite the highly detailed visualization achieved here, 10X/0.25NA chip-MUSICAL outperforms 60X/1.2NA EPI fluorescence in visualizing the membrane structures denoted by the white-dotted oval. **f)** Chip-MUSICAL image acquired with a 60X/1.2NA surpasses the lateral resolution of 60X/1.2NA EPI, reaching down to 194 nm, thus allowing for a sharper visualization of the cellular features previously seen in d).

Further on, to prove the contrast capabilities of the photonic chip-based, we focused our attention on a region of interest within the adenocarcinoma area (yellow box in Figure 3b) and proceeded with fluorescence imaging at higher magnification using four different approaches, namely, epifluorescence microscopy (EPI, Figure 3c), deconvolution microscopy (DV, Figure 3d), photonic chip-based TIRF microscopy (chip-TIRF, Figure 3e), and chip-based FF-SRM (chip-MUSICAL, Figure 3f). Both EPI and deconvolution images were acquired on a commercial DeltaVision Elite High-resolution microscope using a 60X/1.42 oil immersion objective, while the chip-based images were taken with a 60X/1.2 water immersion objective on a custom-made upright microscopy setup (see Supplementary Information S2).

In EPI fluorescence, the off-focus light increases both the foreground and the background signal levels, introducing blur and reducing the contrast ratio between the in-focus signal and its surroundings. This blurring effect can be observed in the magnified views of nuclei and membranes (Figure 3c1 and Figure 3c2, respectively), where the background signal exhibits a characteristic fuzzy appearance inherently present in EPI fluorescence images (Figure 3c3). Hence, to improve the contrast of EPI fluorescence images, a method for out-of-focus signal removal is required.

Deconvolution microscopy (DV) is a well-established diffraction-limited method for contrast enhancement of fluorescent-based images via out-of-focus signal removal^47^. In the context of the colorectal sample used in this study, DV offers a sharper visualization of both the nuclei (Figure 3d1) and the membranes (Figure 3d2), as compared to their corresponding EPI fluorescence images (Figure 3c1 and Figure 3c2, respectively). DV offers high-contrast images near the resolution limit of the microscope (Figure 3d3). However, a close look at Figure 3d1 reveals the overlay of nuclear structures (denoted by a white-dotted box in Figure 3d1), which hampers the interpretation of the data in, for example, nuclei quantification. Arguably, the reason for the nuclear overlay seen in Figure 3d1 is the poor axial resolution. DV microscopy, like any other diffraction-limited technique, has a poorer axial resolution (*d*_*axial*_ = 2λ/*NA*^2^) compared to its lateral resolution (*d*_*lateral*_= λ/2*NA*). In practical terms, for a nuclear marker with emission wavelength λ_*emission*_= 523 nm and an objective lens with numerical aperture *NA* = 1.42, this implies a theoretical axial resolution of ∼520 nm versus a theoretical lateral resolution of ∼184 nm. Hence, the DV method does a good job resolving side-to-side structures, but it struggles to differentiate them when they are too close in the axial direction, as is usually the case with tissue samples.

Photonic chip-based microscopy circumvents this challenge. Besides improving the contrast and the lateral resolution of the fluorescence image^57^, the ultrathin optical sectioning supported by this method enables a clear identification of the nuclei shape (denoted by a white-dotted box in Figure 3e1), which further facilitates the quantification of these structures during histological analysis. In addition, chip-TIRF allows for a sharp visualization of the membrane structures in contact with the propagating waveguide (Figure 3e2). Despite the FFPE colorectal sample being relatively thick (3 µm), the photonic chip-based microscopy offered optical sectioning that resulted in a histological visualization comparable to that obtained through more advanced, complex, and expensive semithin (0.5 µm - 1 µm) and ultrathin (<100 nm) mechanical sectioning of resin-embedded samples used for light and electron microscopy, respectively^58, 59^.

The chip-TIRF image (Figure 3e3) can be further improved by FF-SRM algorithms to achieve an even better visualization of the colorectal sample. Here, by implementing MUSICAL on the chip-TIRF raw data, we substantially improved the contrast over the chip-TIRF images of both nuclei (Figure 3f1) and membranes (Figure 3f2), allowing for a sharp merged image (Figure 3f3) with superior contrast and resolution as compared to all fluorescence-based imaging methods illustrated in Figure 3. Importantly, the chip-based images (TIRF and MUSICAL), were acquired with a lower numerical aperture objective (*NA* = 1.2) compared to the lens (*NA* = 1.42) used for the EPI-based imaging (EPI and DV). This explains, for example, the superior boundary sharpness obtained in the DV image of nuclei (Figure 3d1) as compared to the corresponding chip-MUSICAL image (Figure 3f1). Nevertheless, photonic chip-based microscopy outperforms EPI-based methods offering superior contrast visualization of FFPE samples over large fields of view. In the next section, we explore the lateral resolution scalability offered by photonic chip-based microscopy for the study of paraffin-embedded sections.

### Super-resolution chip-based histology over scalable fields of view

Histological examination, regardless of the labeling method, often requires visualization of the tissue sample with diverse levels of magnification. Typically, low-magnification objectives are used for contextual interpretation of the sample, while higher magnification lenses are employed for the study of finer details and subcellular structures within a region of interest for the observer. Low magnification, for example, assists in the identification of large morphological features in the tissue such as cellular epithelium, glands, vessels, inflammation processes, and cancer proliferation, whereas higher magnification enables detailed views of cellular features such as organelle dynamics, nuclei shape, extracellular matrix, and protein localization. In this part of the study, we used an FFPE prostate cancer sample to illustrate the enhanced resolution offered by photonic chip-based microscopy over scalable fields of view.

Here, a prostate specimen was taken from a patient, fixed, grossed, processed, and further embedded in paraffin following standard FFPE method. Thereafter, the tissue block was sectioned to a 3 µm thin slice using a microtome and floated on a water bath. Then, the sample was scooped onto a photonic chip and further stained for nuclei and membranes (see detailed preparation protocols in the Materials and Methods section).

The fluorescence imaging was performed on a custom-made microscopy setup equipped with a photonic chip module and epifluorescence (see Supplementary Information S2). Along with the chip-based images, EPI images were also collected for the sake of direct comparison. First, the tumor area was located using a 10X/0.25NA objective lens. Thereafter, a region of interest was further examined using a 60X/1.2NA water immersion objective lens. Upon collecting individual channels, the images were pseudo-colored (nuclei in cyan, and membranes in magenta), and further merged. Figure 4a illustrates a 10X EPI fluorescent multicolor image of the prostate cancer sample with the presence of multiple glands forming cancer cells. Notably, the vignetting observed in the image, namely, the darker areas toward the borders of Figure 4a, originates from the gaussian distribution of the EPI illumination. This non-uniformity not only introduces grid-like stitching artifacts as shown in Figure 3b, but also constrains the exploitable field of view to the center of the EPI image, thereby decreasing the imaging throughput and quantitative capabilities of the microscopy system. This is similar to other approaches that have been proposed to achieve a flat field EPI illumination^60–63^, which require additional optical components and calibration.

Photonic chip-based microscopy circumvents this issue providing the sample with a uniform illumination via waveguide light propagation. Figure 4b shows a 10X chip-TIRF multicolor image of the same sample region as in Figure 4a. Contrary to EPI, chip-TIRF enables a high-contrast observation of the prostate tissue sample across the whole field of view (see Supplementary Video V4). To acquire the chip-based image in Figure 4b, the excitation laser was sequentially coupled to each one of the waveguides in the field of view while TIRF images were collected with a low-magnification and low numerical aperture (10X/0.25NA) objective lens. Upon FF-SRM post-processing via MUSICAL, the individual images were finally merged to obtain a multicolor high-contrast image across a field of view spanning 1.3 mm × 1.3 mm (see Supplementary Information S8). This, in our opinion, is an important advantage offered by photonic chip-based microscopy for histology. By decoupling the excitation and the collection light paths, it is possible to uniformly excite large sample areas through evanescent field illumination, and capture the fluorescent emission essentially with any microscope objective, regardless of its numerical aperture and magnification. Conventional TIRF microscopy techniques, on the contrary, require dedicated EPI objective lenses with high numerical aperture and high magnification (for example, 60X/1.49NA) that, due to the non-uniform Gaussian illumination distribution, support an exploitable field of view of approximately 50 µm × 50 µm (ref. 21). Certainly, the waveguide spacing gaps of 25 µm (dark horizontal lines in Figure 4b) hinder the full visualization of the sample features inside the image field of view. However, this is not a scientific limitation. Future chip designs can address this issue by, for example, widening the waveguides, reducing the spacing gap down to 1 µm, or simply using chips with slab geometries with no gaps.

Further on, a zoom-in view of the white-dotted box areas in Figure 4a and Figure 4b, respectively, revealed the advantages of photonic chip-based microscopy (Figure 4d) over conventional EPI fluorescence microscopy (Figure 4c). In particular, chip-MUSICAL not only improved the sample contrast but also allowed the visualization of membrane structures (denoted by the white-dotted oval inside Figure 4d) that otherwise were not visible in EPI (denoted by the white-dotted oval inside Figure 4c). To estimate the resolution of these two microscopy methods, we performed a decorrelation analysis^64^ over the 10X images of membranes (see Supplementary Information S9). While the EPI fluorescence image rendered a lateral resolution of 1.56 µm, chip-MUSICAL delivered nearly a 1.2-fold improvement compared to EPI, achieving a lateral resolution of 1.31 µm.

To achieve an even higher resolution over the same region as in Figure 4c and Figure 4d, we swapped the collection objective lens to a higher magnification and higher numerical aperture (60X/1.2NA) and repeated the image acquisition steps both for EPI fluorescence and chip-MUSICAL. While the 60X images provided a clearer definition of the cellular elements as compared to its corresponding 10X images, a decorrelation analysis (see Supplementary Information S10) revealed a ∼2.4-fold lateral resolution gap between the high-magnification EPI fluorescence (Figure 4e) and the high-magnification chip-MUSICAL (Figure 4f). More specifically, 461 nm in EPI versus 194 nm in chip-MUSICAL using a 60X/1.2NA objective lens. Notably, the 10X/0.25NA chip-MUSICAL (Figure 4d) outperformed the 60X/1.2NA EPI fluorescence (Figure 4e) in, for example, visualizing the membrane features located inside the white-dotted oval. This result suggests that chip-based histology could potentially enable high-resolution and high-throughput imaging of tissues over large fields of view, employing low magnification objectives, which could further facilitate the screening of FFPE samples in clinical settings.

### Chip-based CLEM histology

Both fluorescence and electron microscopy provide unique information about the samples under observation. While fluorescence microscopy offers high specificity and high-contrast details, electron microscopy enables the visualization of features down to the ultrastructural level. Hence, combining these two microscopy modalities is advantageous for histological analysis. While several correlative light and electron microscopy (CLEM) studies exist in the histological field ^65–67^, the tissue preservation method employed in conventional approaches consisted mainly of cryo-preservation or resin embedding. Over the last decade, however, innovative research has shown the potential of incorporating FFPE samples into the CLEM analysis to, for example, study inflammatory processes^68^, identify virus particles in diseased organs^69^, and render a high-definition topographic visualization of thick tissue sections^70^. Here, we propose photonic chip-based microscopy as a feasible platform for combining high-contrast fluorescence-based imaging with high-resolution scanning electron microscopy (SEM) of paraffin-embedded sections.

In this part of the study, we used a human placental tissue section to demonstrate the compatibility of the photonic chip for correlative light and electron microscopy of FFPE samples. Here, a chorionic villi sample was taken from the fetal side of a human placenta, dissected, and further embedded in paraffin following standard FFPE method. Thereafter, the tissue block was sectioned into a 3 µm slice and fluorescently labeled for nuclei and membranes (see detailed preparation protocols in the Materials and Methods section).

A common challenge for correlative imaging is finding the same region of interest across different microscopy systems. The photonic chip can be fabricated with a landmark coordinate system along the spacing gaps between the waveguides that facilitates the navigation through the sample^71^. To perform CLEM imaging on the FFPE placental sample, we first used a low-magnification objective (10X/0.25NA) in EPI fluorescence mode to find a region of interest (ROI). Thereafter, we switched to bright field imaging mode to visualize the landmarks around the ROI and, further on, we combined the bright field and the EPI images (Figure 5a) to obtain a combined view of both the landmarks (Figure 5c) and the fluorescent signal (membranes in magenta and nuclei in cyan) within the same field of view. Then, to achieve a detailed visualization of the ROI, we transitioned to a higher magnification collection objective (60x/1.2NA) and performed chip-based FF-SRM via MUSICAL (Figure 5e). Upon completion of fluorescence imaging, the coverslip was carefully removed and the sample was subsequently dehydrated and coated with a gold/palladium alloy following the preparation steps for SEM (see Materials and Methods section). Thereafter, the chip was placed on an SEM device (GeminiSEM 300, Zeiss) and imaged at low magnification to navigate through the chip landmark coordinates (Figure 5d) and quickly find the region of interest (Figure 5b). Further SEM magnification allowed for a topographic visualization (Figure 5f) of the same ROI obtained via waveguide illumination (Figure 5e). This feature enabled a high-detail identification of structures observed both in fluorescence and in electron microscopy. For example, a zoomed-in view of the placental tissue in Figure 5g revealed the microvilli (MV) brush border outlining the apical side of the syncytiotrophoblasts (SYN). Similarly, a zoomed-in view of the same region in SEM (Figure 5h) validated the previous observation indicating that, in fact, the microvilli imaged through chip-based illumination matched the structures at the bottom of the tissue sample in direct contact with the waveguide surface. This demonstrated the ground truth for on-chip MUSICAL with SEM images. Moreover, the overlay between the chip-based and the SEM images (Figure 5i) showed a perfect correlation between the two imaging methods, which was further scalable to the complete field of view (see Supplementary Video V5).

**Figure 5.**
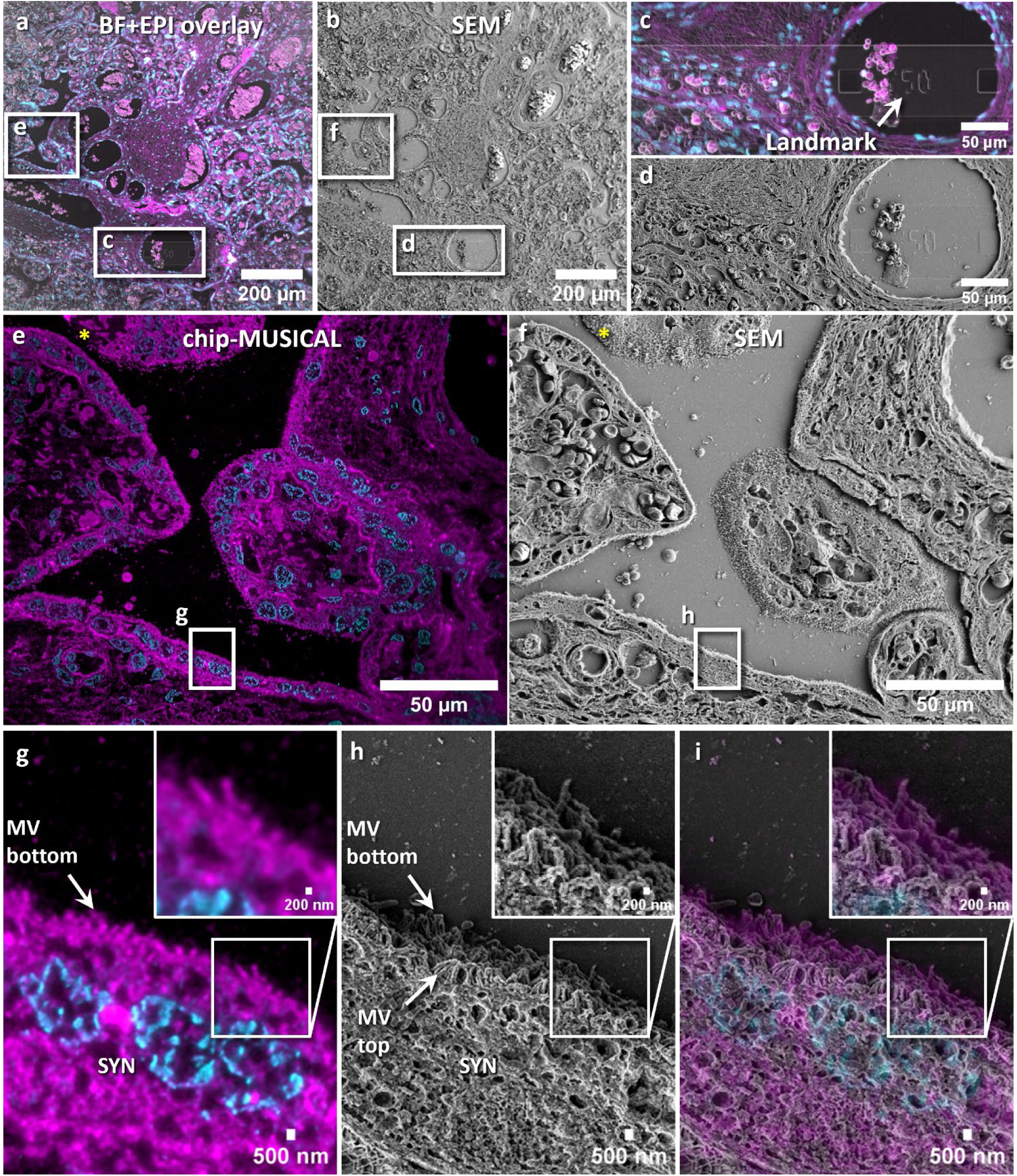
Chip-based CLEM histology of an FFPE placental tissue section. **a)** 10X magnification bright field and EPI fluorescence overlay image of a region of interest within the placental sample. Membranes are shown in magenta and nuclei in cyan. The white box at the bottom denotes the location of a landmark that is used to find the sample in the scanning electron microscope (SEM). The white box to the left side of the image denotes the region of interest imaged at higher magnification in e). **b)** Low-magnification SEM image of the same region imaged in a). The white box at the bottom denotes the location of the landmark coordinate described in a). The white box to the left side of the image denotes the region of interest imaged at higher magnification in f). **c-d)** Zoom-in view of the landmark coordinate (Y50) used for the localization of the same region of interest among different microscope systems. **e)** Chip-based super-resolution image of chorionic villi reconstructed via MUSICAL. The white box at the bottom of the image denotes the location of a region of interest further described in g). The yellow asterisk denotes a detached sample region (see details in Supplementary Information S12). **f)** SEM image of the same sample region imaged in e). The white box at the bottom of the image denotes the location of a region of interest further described in h). The yellow asterisk denotes a detached sample region (see details in Supplementary Information S12). **g)** Zoom-in view of the chip-based image reveals the microvilli brush border outlining the syncytiotrophoblast (SYN). **h)** Zoom-in view of the same sample region as g) confirms the presence of microvilli along the volume of the sample. **i)** The image overlay shows an excellent correlation between the chip-based and the SEM images. Importantly, the microvilli outline in g) matches the location of the structures in direct contact with the waveguide surface at the bottom of h).

The CLEM study also allowed us to further evaluate our hypothesis of FFPE sample detachment. In the case of the placental section (see details in Supplementary Information S12), for example, the fluorescence signal discontinuity observed in the chip-TIRF image around the apical side of the upper villus (yellow asterisk in Figure 5e), matched the location of a micro-detachment gap visualized in the subsequent SEM image (yellow asterisk in Figure 5f). Supplementary Information S13 summarizes the optimization steps carried out to enable optimal sample adhesion.

## Conclusion and Discussion

The FFPE on-chip method proposed here offers advantages for the practical adoption of fluorescence-based super-resolution histology (see details in Supplementary Information S11). In particular, the ultrathin optical sectioning combined with the MMI illumination pattern modulation supported by the photonic chip allows for a seamless implementation of FF-SRM methods such as MUSICAL, which otherwise fail for densely labeled samples such as FFPE tissue sections, as shown in Figure 2. Moreover, the TIRF illumination offered by the photonic chip enables high-contrast images, allowing for accurate identification of morphological features irrespective of the objective lens magnification used, as highlighted in Figure 4. The photonic-chip illuminates the sample along the entire length of the waveguide and, with future integration of microlens arrays^72^ in the collection light path, it could be envisioned a dramatic speed enhancement of the photonic chip-based system, supporting image acquisition along the entire length of guiding waveguides. Similarly, the incorporation of multiplex coupling automation could also boost the imaging throughput of photonic chip-based microscopy by, for example, allowing for simultaneous image acquisition of several waveguides in a similar fashion to existing whole slide scanners used in routine histopathology laboratories. Another key advantage of photonic chip-based microscopy is the uniform sample visualization delivered by waveguide illumination (Figure 4). Chip-TIRF enables the exploration of the whole image field of view, therefore increasing the imaging throughput as compared to conventional EPI-based microscopy methods that suffer from reduced field flatness. We envision that, upon further assistance of machine-learning approaches^49, 73^, the big data of high-resolution images over large fields of view supported by photonic chip-based microscopy could also facilitate histological interpretability and diagnosis.

Another benefit of the photonic chip for histology is its compatibility with the conventional FFPE sample preparation protocols. Notably, the chips used in this study resisted all the harsh processing steps associated with deparaffinization, rehydration, and antigen retrieval for fluorescence (immuno)labeling. Admittedly, the relatively small size of the photonic chips utilized here (roughly, 2.5 cm × 2.5 cm), resulted in manual processing of the samples (see Supplementary Information S5). However, the chips could be manufactured with the same dimensions as traditional glass slides, enabling compatibility with commercial histochemical processing devices and fluorescence immunostaining machines. Similarly, the current imaging limitation imposed by the spacing gaps (Figure 4b) could easily be addressed by redesigning the photonic chip top layer geometry to accommodate wider optical waveguides, for example, ∼5 mm wide, which can fill the entire field of view of low magnification objectives, or by narrowing the spacing gap width down to ∼1 µm or using a slab waveguide with no gap.

In this study, we focused our florescence labeling efforts on two direct markers, for nuclei and membranes, respectively, to enable a one-to-one comparison with standard H&E staining (see details in the Materials and Methods section). However, photonic chip-based microscopy is also compatible with fluorescence immunolabeling approaches (Supplementary Information S6), enabling a highly specific visualization of histological features that are relevant both for research and clinical diagnosis. We envision that, upon integration of microfluidic systems, the photonic chip could also be used in combination with advanced fluorescent labeling techniques such as DNA-PAINT^74^ and exchange-PAINT^75^ for high-content multi-omic screening^76^, as well as for multiplex super-resolution imaging of FFPE samples via SMLM^3^.

We did encounter sample detachment challenges that deserve further attention, as they compromise the imaging capabilities of the photonic chip (see Supplementary Information S12). Future studies should address the adhesion of the paraffin-embedded slices to the chip surface, to successfully achieve whole-sample TIRF imaging. Alternative histochemical processing methods such as resin embedding could also be explored to enable optimal sample attachment and subsequent chip-based imaging.

Importantly, chip-TIRF is limited to a 2D visualization of the part of the sample in contact with the waveguide surface. However, as demonstrated here, the photonic chip is compatible with alternative fluorescence-based imaging modalities, such as EPI and DV, which provide a complementary volumetric view of the tissue section that further assists in the histological interpretation. In addition, the embedded landmark coordinate system allows for effortless navigation over large tissue samples to localize the same regions of interest among different imaging systems. This proved useful, for example, in the correlative light and electron microscopy study of the placental section. Lastly, the combination of section thickness (2 – 4 µm) and refractive index heterogeneity imposed by the FFPE samples, can introduce optical aberrations in the form of light scattering^77^ that could potentially hinder the contrast and resolution of the chip-based microscopy. To mitigate this, two approaches can be explored: a) homogenizing the refractive index of the tissue via optical clearing^78^, or b) using transparent chips in combination with an inverted microscope setup to avoid light scattering of the fluorescent signal^57^.

## Acknowledgments

The authors are grateful to the histo-technicians who performed the tissue sectioning for this study. These include Mona Pedersen and Premasany Kanapathippillai from UiT – The Arctic University of Norway (Tromsø, Norway), and Marna Lill Kjæreng from the Institute for Cancer Genetics and Informatics at Radium Hospital (Oslo, Norway). The authors would like to thank Dr. Azeem Ahmad for assisting with the integration and system development of the photonic chip-based setup. The authors would like to express their gratitude to Prof. Malgorzata Lekka at the Institute of Nuclear Physics, Polish Academy of Sciences (Krakow, Poland), for facilitating the AFM measurements of the FFPE sections. The authors would like to thank Dr. Seshi Sompuram and Dr. Steve Bogen from Boston Cell Standards (Boston, USA) for their support with the PI coating test of the chips. The authors would like to acknowledge Randi Olsen at the Advanced Microscopy Core Facility at UiT – The Arctic University of Norway (Tromsø, Norway) for her assistance in preparing the placental sample for SEM. The authors appreciate the support of Dr. Firehun T. Dullo (SINTEF MiNaLab, Oslo, Norway), for the design and fabrication of the Si_3_N_4_ photonic chips. BSA acknowledges the funding from the Research Council of Norway, projects # NANO 2021–288565 and # BIOTEK 2021–285571, and from the European Innovation Council (EIC), EIC-Transition project # 101058016. KA acknowledges funding from the Research Council of Norway project # 288082 (FRIPRO YOUNG program).

## Author contribution

BSA conceived the idea, secured the project, and supervised the project. LEVH and BSA planned and coordinated the experiments. LEVH, VD, and HM performed sample labeling, chip-TIRF imaging, and post-processing of the data. VD built the chip-based microscope setup. SA and KA assisted with MUSICAL reconstructions. HED coordinated and provided the colorectal and prostatic FFPE sections. MP assisted in the interpretation of the colorectal and prostatic samples. HM and LEVH performed the SEM imaging of the placental section. JCT assisted in the photonic chip design and fabrication logistics and the design and assembly of the chip-based microscope setup. DHH assisted with the software integration and mechanical engineering for chip-TIRF imaging. GA provided the placental sample. MN helped with the collection, preservation, and image interpretation of the placental sample. KAF collected and preserved the mouse kidney tissue (supplementary) and assisted with the renal image interpretations. BZ assisted with the measurements and interpretation of the AFM data of the colorectal sample (supplementary). LEVH analyzed the data, prepared the figures and the supplementary videos of the manuscript, and wrote the manuscript. LEVH and BSA wrote the first draft of the manuscript and all authors contributed to writing and revising selected sections of the manuscript.

## Materials and Methods

### Photonic chip description and fabrication

The photonic chip is made of three layers (see Figure 1a), namely, a substrate of silicon (Si), an intermediate layer of silicon dioxide (SiO_2_), and a top optical waveguide layer of silicon nitride (Si_3_N_4_). The high refractive index contrast between the waveguide material (*n* ≈ 2.0) and the sample medium (n ≈ 1.4), allows a confined propagation of the excitation light along the waveguide via total internal reflection (TIR), which in turn provides evanescent field excitation to the fluorescent molecules in the vicinity of the waveguide (see Supplementary Information S4), hence enabling chip-based total internal reflection fluorescence (chip-TIRF) microscopy. Previous studies have explored diverse waveguide geometries for chip-TIRF microscopy. These include slab, rib, and strip waveguides^25^. In this work, we chose 140 nm height uncladded strip waveguides with widths varying from 200 µm to 1000 µm (see Figure 1b).

The photonic chips used here were manufactured at SINTEF MiNaLab (Oslo, Norway), following a standard photolithography CMOS fabrication process as described elsewhere^25, 26^. Briefly, a silicon dioxide layer of 2 µm thickness was thermally grown on a silicon chip. Thereafter, a layer of silicon nitride was deposited using low-pressure chemical vapor deposition at 800°C. To delineate the diverse strip waveguide geometries, reactive ion etching (RIE) over a photoresist mask was employed. Next, the remaining photoresist was removed before depositing a top cladding layer of 2 µm via plasma-enhanced chemical vapor deposition at 300°C. To allow for TIRF imaging, the top cladding was removed from the central portion of the chip using RIE and wet etching. All waveguides were fabricated with a thickness of 140 nm.

### Sample origin and ethical approvals

Human prostate samples were obtained from male patients following radical prostatectomy at the Norwegian Radium Hospital (Oslo, Norway). Colorectal samples were obtained after resection from human patients at Aker University Hospital (Oslo, Norway). Full-term placentas from healthy patients were collected immediately after delivery at the University Hospital of North Norway (Tromsø, Norway). All samples were fixed in formalin and embedded in paraffin following standard histological methods. All samples were anonymized following the guidelines of the Regional Committees for Medical and Health Research Ethics of Norway (REK). No personal data was attached to the biological samples used in this study.

### FFPE tissue sectioning

The paraffin blocks were cooled down at 4 °C before sectioning into 3 µm to 4 µm slices. The FFPE samples were sectioned using two automatic microtomes (HM 355S, Thermo Fisher Scientific) equipped with a water bath. The FFPE prostate and colorectal samples were sectioned at the Institute for Cancer Genetics and Informatics (Oslo, Norway). The placental samples were sectioned at the Department of Clinical Medicine at UiT – The Arctic University of Norway (Tromsø, Norway).

### H&E staining and widefield imaging of the colorectal sample

Upon sectioning, the paraffin-embedded colorectal slice was placed on a superfrost plus glass slide (J1800AMNZ, Epredia) and histochemically stained with H&E. Thereafter, the colorectal sample was imaged on a whole slide scanner device (Virtual Slide System V120, Olympus) using a 20X/0.85NA objective lens in widefield illumination. The individual images were automatically stitched in a tile-mosaic array by the proprietary software available on the scanning device. An experienced pathologist identified and annotated the four regions of the sample (adenocarcinoma, benign epithelium, necrosis, and smooth muscle).

### On-chip FFPE fluorescence labeling

The on-chip FFPE samples were manually processed as follows (see Supplementary Information S5): the photonic chips were first coated with 0.1 % poly-L-lysine (P8920, Sigma-Aldrich) and then glued to conventional glass slides for easier handling using picodent twinsil dental glue (1300-1000, Picodent). Upon sectioning, the paraffin-embedded slices were scooped from the water bath and deposited on the central portion of the photonic chips (see Supplementary Video V3). The sections were dried on a flat surface at room temperature (1 × 1 h) and then transferred to a 60 °C incubation oven (TS4057, Termaks) for overnight melting (approx. 16h) of the paraffin. Thereafter, the samples were transferred to a wash-N-dry coverslip rack (Z688568, Sigma-Aldrich) for further processing steps in glass beakers of 100 mL capacity. The samples were deparaffinized in xylene (3 × 5 min), and rehydrated in descendent series of ethanol starting from 100 % (2 × 10 min), to 96 % (2 × 10 min), and finally 70 % (1 × 10 min). Subsequently, the samples were washed with MilliQ water (5 min) before placing them on a flat surface for fluorescence labeling. For membrane staining, the samples were incubated (1 × 15 min) in a 1:2000 solution of MitoTracker Deep Red FM (M22426, Invitrogen) in phosphate-buffered saline (PBS), (D8662, Sigma-Aldrich). Except for the FFPE placental section in Figure 2, all the other FFPE sections in this study were subsequently labeled for nuclei. For this, the samples were rinsed (1 × 10 sec) and washed with PBS (2 × 5 min), and further incubated in a 5 µM solution of Sytox Green (S7020, Invitrogen) in PBS for nuclei staining. Next, the samples were rinsed (1 × 10 sec) and washed with PBS (2 × 5 min) before mounting with glycerol (G5516, Sigma Life Science) and covering with #1.5 coverslips of 22 mm × 22 mm (631-0125, VWR). The staining and washing steps were performed with single-channel micropipettes (Finnpipette F2, Thermo Scientific) and an aspirator pump (FTA-1, BioSan). The incubation volumes were between 300 µL and 500 µL depending on the sample area dimensions. The samples were sealed with picodent twinsil dental glue and stored at 4 °C protected from the light until imaging. The microscopy observations were performed within 1 to 3 days after labeling. Supplementary Information S16 provides a detailed description of the materials and reagents used for on-chip FFPE fluorescence labeling.

### EPI fluorescence and DV imaging of the FFPE colorectal sample

Upon on-chip deposition and fluorescence labeling, the photonic chip containing the colorectal cancer was glued to a standard glass slide employing adhesive tape and turned upside down for observation on an inverted fluorescence microscope (DeltaVision Elite Deconvolution Microscope, GE Healthcare). To visualize the totality of the colorectal sample (Figure 3b), a 10X/0.4NA air objective lens in EPI fluorescence mode was used. To achieve multicolor images, two consecutive images were taken for each field of view. The far-red channel (λ_excitation_= 632 ± 11 nm, λ_emission_ = 679 ± 17 nm) for membranes, and green channel (λ_excitation_ = 475 ± 14 nm, λ_emission_ = 525 ± 24 nm) for nuclei. The exposure time and the illumination power were adjusted to obtain a maximum of 10000 grayscale counts on the camera chip. The frames were automatically stitched into an 8 × 8 tile mosaic image by the built-in software package SoftWoRx available on the instrument and further pseudo-colored in the open-source software Fiji^79^. The regions of interest were mapped to the annotations made by the expert pathologist on the H&E colorectal slide.

For a higher magnification view of the colorectal sample, a 60X/1.42NA oil immersion objective was used. To obtain optimal results, we used an immersion oil with a refractive index *n* = 1.516. For EPI fluorescence observation (Figure 3c), two consecutive single-plane images were taken over the same field of view, following the imaging strategy of 10X EPI fluorescence. For DV imaging (Figure 3d), a total of 48 z-plane frames with z-steps of 250 nm were collected on each imaging channel along the optical axis of the objective. Thereafter, individual z-planes were deconvolved with the built-in SoftWoRx package. Finally, a deconvolved frame at the center of the stack (z = 29) was chosen for the analysis.

### Chip-based imaging of FFPE samples

To build up the photonic chip-based microscope setup, a modular upright microscope (BXFM, Olympus) and a custom-built photonic chip module were used (Supplementary Information S2). The excitation light was provided by a multi-wavelength fiber-coupled laser (iChrome CLE, Toptica), which was expanded and collimated through an optical fiber collimator (F280APC-A, Thorlabs) to fill the back aperture of the coupling objective (NPlan 50X/0.5NA, Olympus). The excitation wavelengths used in this study were mainly λ_1_ = 640 nm, and λ_2_ = 488 nm. To optimize the light coupling and subsequent scanning along the waveguide input facet, both the optical fiber collimator and the coupling objective were mounted onto an ensembled translation system consisting of a miniature piezo-controllable X-axis stage (Q-522 Q-motion, PI) fitted onto an XYZ translation stage (Nanomax300, Thorlabs). The photonic chips were placed on a custom-made vacuum chuck fitted on an X-axis translation stage (XRN25P, Thorlabs) for large-range scanning of parallel waveguides. Upon coupling the excitation light onto a selected waveguide, the fluorescent emission of the samples was accomplished via evanescent field excitation (Figure 1a). The fluorescent signal was collected using a variety of MO lenses depending on the desired FOV, magnification, and resolution. These include 4X/0.1NA air, 10X/0.25NA air, 20X/0.75NA air, and 60X/1.2NA water immersion. To block out the excitation signal at each wavelength channel, an emission filter set comprising a long-pass filter and a band-pass filter was employed (details in Supplementary Information S2). After passing the 1X tube lens (U-TV1X-2, Olympus), the fluorescence signal was captured by an sCMOS camera (Orca-flash4.0, Hamamatsu). Both the camera exposure time and the laser intensity were adjusted according to the experimental requirements. In the case of chip-TIRF imaging, the camera exposure time was set between 50 ms and 100 ms, and the input power was gradually increased until the mean histogram values exceeded 500 counts. Typical input powers were between 10 % and 60 % depending on the coupling efficiency and the waveguide width. To minimize photobleaching of the fluorescent tags, the chip-TIRF acquisition was sequentially performed from less energetic (λ_1_ = 640 nm) to more energetic (λ_2_ = 488 nm) excitation wavelengths. To achieve uniform illumination of the sample, the coupling objective was scanned along the waveguide’s input facet width in lateral steps of < 1 µm while individual images were acquired. Usually, image stacks between 200 and 1000 frames were collected. To obtain multicolor images, the process was repeated at different excitation wavelengths according to the fluorescent markers available on the sample. To ensure mechanical stability, the microscope body was fixed to the optical table, while the photonic chip module was placed onto a motorized stage (8MTF, Standa) for scanning across the XY directions. An optical table (CleanTop, TMC) was used as the main platform for the chip-TIRFM setup.

### EPI fluorescence imaging of FFPE samples on a chip-based setup

To enable a volumetric view of the FFPE samples in bright field (BF) and EPI fluorescence modes, respectively, the chip-based microscope setup was equipped with a halogen lamp (KL1600 LED, Olympus) and a secondary multi-wavelength fiber-coupled laser (iChrome CLE, Toptica) with the same laser spectrum as for the waveguide illumination. Both the BF and the EPI signals were acquired using a beam splitter and a series of dichroic mirrors (details in Supplementary Information S2).

For the FF-SRM comparison shown in Figure 2, the placental section was imaged in EPI fluorescence mode, using a 640 nm laser wavelength at 3 % transmission power. To collect the intrinsic fluorescence fluctuations of the fluorophores, the camera was set to a short acquisition time of 10 ms. The collected EPI image stack consisted of 500 frames. For illustration purposes, the image stack shown in Supplementary Video V2 was shortened to only the first 200 frames.

### Correlative light-electron microscopy of placenta

After chip-TIRF imaging, the placental sample was brought to the Advanced Microscopy Core Facility at UiT – The Arctic University of Norway (Tromsø, Norway) for further preparation and SEM imaging. First, the picodent twinsil glue was removed from the edges of the coverslip. Thereafter, the photonic chip was immersed and washed with PHEM buffer^46^ to dilute the glycerol and facilitate the coverslip detachment. Then, the sample was treated in freshly made 1 % tannic acid in 0.15 M cacodylic buffer for 1 hour, followed by 1 % osmium tetroxide (OsO_4_) in 0.1 M cacodylic buffer for 1 hour, and dehydrated in incremental ethanol series (30 %, 60 %, 90 % for 5 minutes each, and 5 times 100 % ethanol for 4 minutes). Next, the sample was incubated twice in hexamethyldisilazane for 2 minutes each and carbon-taped to a 25 mm diameter SEM stub followed by silver glue for electrical conduction. Then, the chip with the sample was stored in a desiccator overnight to allow dehydration. Subsequently, the chip was coated with a 10 nm layer of gold/palladium alloy and brought to a scanning electron microscope (GeminiSEM 300, Zeiss) for SEM imaging at low accelerating voltage using an in-lens detector. Also, to enable a topographic view of the placenta, the sample was tilted 25 degrees and imaged with a secondary electron detector using low accelerating voltage.

### Image processing and analysis

The acquired chip-TIRF stacks were computationally averaged using the *Z Project* tool in Fiji^79^. Thereafter, the averaged channels were merged and pseudo-colored with the *Merge Channels* tool (FIJI) to obtain multicolor chip-TIRF images.

For chip-MUSICAL reconstructions, a Python implementation of soft-MUSICAL^80^ was used. The size of the sliding window was set to 7 × 7 pixels, and the sub-pixelation to 5. Pixel sizes of 650 nm and 108 nm were used for the 10X/0.25NA and the 60X/1.2NA image stacks, respectively. The signal wavelength was set according to the emission wavelength of the given marker, namely 670 nm for the MTDR, and 520 nm for the Sytox Green. The MUSICAL reconstructions were further adjusted using the *Log* transform and pseudo-colored in FIJI.

For CLEM, the acquired chip-TIRF stacks were first processed with soft-MUSICAL and then correlated with the SEM image using the TrakEM2 plugin^81^.

### Figure and video rendering

The schematic representations shown in this manuscript were created with BioRender.com. The supplementary videos were assembled with Microsoft’s Clipchamp app.

## Supplementary Information

### S1. Mode-averaging for chip-TIRF

In this study, we used photonic chips with strip waveguides of various widths (200 µm, 400 µm, 600 µm, and 1000 µm) to evaluate the performance of chip-TIRF under different geometrical configurations. For chip-TIRF imaging, a side illumination laser is coupled onto a selected waveguide using a microscope objective. Upon coupling, the excitation light propagates through the waveguide in the form of an anisotropic intensity distribution called multi-mode interference (MMI) pattern, which can be further modulated by changing the position of the coupling objective (Supplementary Figure S1a). To attain uniform illumination of the specimen, the coupling objective is scanned along the input facet of the chip while individual frames are acquired (Supplementary Figure S1b). To achieve a uniform chip-TIRF image, the collected image stack is then averaged (Supplementary Figure S1c). Supplementary Video V1 provides a visual animation of the on-chip mode averaging process.

**Supplementary Figure S1.**
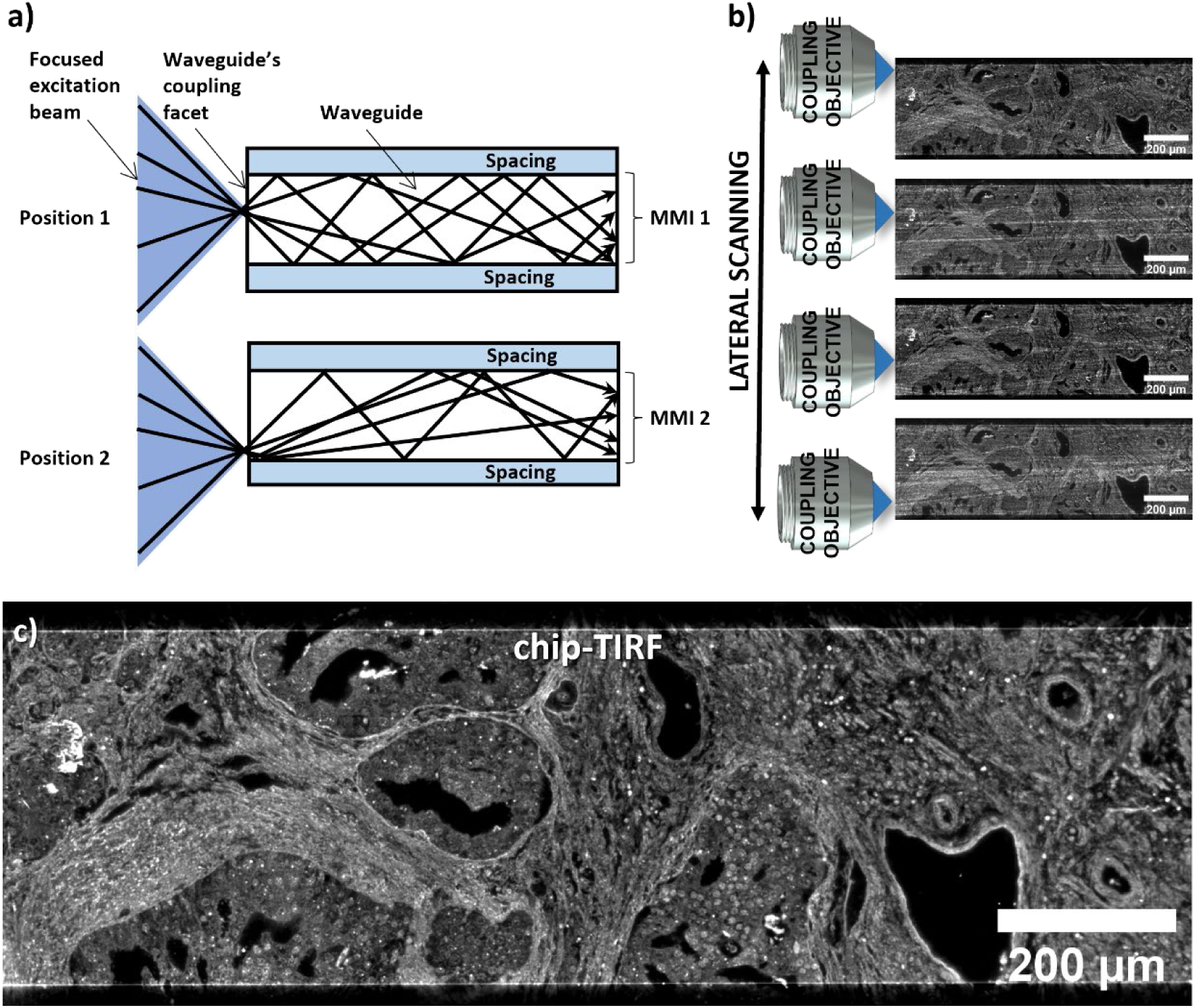
Mode-averaging for chip-TIRF images. **a)** Upon coupling the excitation laser, the excitation light propagates along the selected waveguide via total internal reflection. The geometry of the waveguide supports multi-mode interference (MMI) patterns that can be modulated by scanning the coupling objective relative to the input facet of the chip. **b)** An image stack is collected by acquiring one frame for each position of the coupling objective. **c)** The image stack is then averaged in FIJI to obtain a diffraction-limited high-contrast chip-TIRF image.

### S2. Chip-based microscopy setup

The chip-based microscope is composed of two main sections, namely the collection module and a photonic chip module, as illustrated in Supplementary Figure S2a. The collection module consists of a commercial upright microscope equipped with an emission filter set (see Supplementary Table S2), an sCMOS camera, and conventional microscope objective lenses of diverse magnifications, which can be interchanged depending on the imaging needs. The photonic chip module consists of a set of translation stages to enable precise control of the excitation laser source for chip-based TIRF imaging. By using a beam splitter and dichroic mirrors, it is also possible to image the sample via widefield and EPI fluorescence illumination. Supplementary Figure S2b and Supplementary Figure S2c provide a detailed view of the chip-based microscope setup.

**Supplementary Table S2.** Dichroic mirrors and bandpass filters used for EPI and chip-TIRF image acquisition.

| Excitation wavelength (nm) | Emission filter set |  |
| --- | --- | --- |
|  | Dichroic mirror/Long-pass filter (nm) | Band-pass filter (nm) |
| 488 | 515 (CHR-T515LP, Chroma) | 535/30 (CHR-ET535/30M, Chroma) |
| 561 | 585 (CHR-T585LPXR, Chroma) | 630/75 (CHR-ET630/75M, Chroma) |
| 640 | 655 (CHR-AT655DC, Chroma) | 690/50 (CHR-AT690/50M, Chroma) |

**Supplementary Figure S2.**
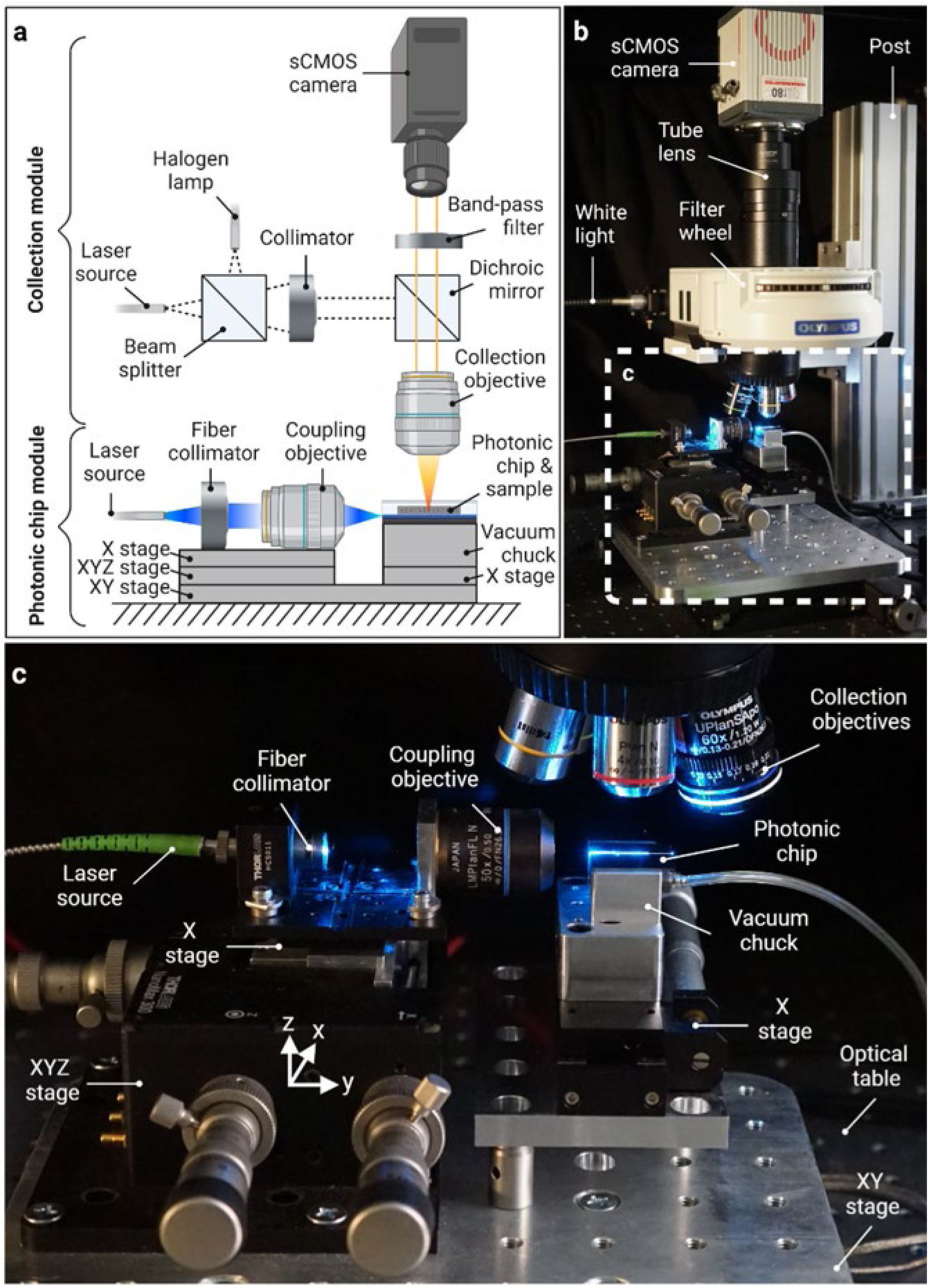
Chip-based microscopy setup. **a)** Schematic representation of the chip-based microscopy setup illustrating the collection and the photonic chip modules, respectively. **b)** Side view of a chip-based microscope setup. The white-dotted box denotes the photonic chip module shown in c). **c)** Magnified view of the photonic chip module including part of the collection module (collection objectives).

### S3. Depth of field of microscope objectives

The depth of field (DOF) refers to the distance between the closest and the further objects that can be sharply imaged along the focal plane of a lens. In microscopy, the DOF is dictated by the formula DOF= λ*n*⁄*NA*^2^, where λ is the wavelength of the excitation light, *n* is the refractive index of the medium between the coverslip and the microscope objective, and NA is the numerical aperture of the microscope objective. The DOF plays a crucial role both in the lateral resolution and contrast of fluorescence microscopy. Supplementary Figure S3 illustrates a theoretical simulation of DOF vs the NA of commonly available air (*n* = 1), water immersion (*n* = 1.33) and oil immersion microscope objectives using the aforementioned formula. As shown in the graph, low NA objectives exhibit larger DOF while higher NA objectives allow for shorter DOF. In EPI fluorescence, low NA objectives are usually employed for a contextual view of the sample (for example, in Figure 4a) whereas, for detailed views, higher NA objectives are needed. However, the DOF of high NA lenses still limits the axial resolution of advanced microscopy techniques such as DV (Figure 3d) and FF-SRM methods such as MUSICAL (Figure 2a). Photonic chip-based microscopy, on the contrary, supports <50 nm optical sectioning via evanescent field waveguide-based excitation (see Supplementary Information S4), allowing for high-contrast fluorescent imaging through virtually any microscope objective, irrespective of its DOF.

**Supplementary Figure S3.**
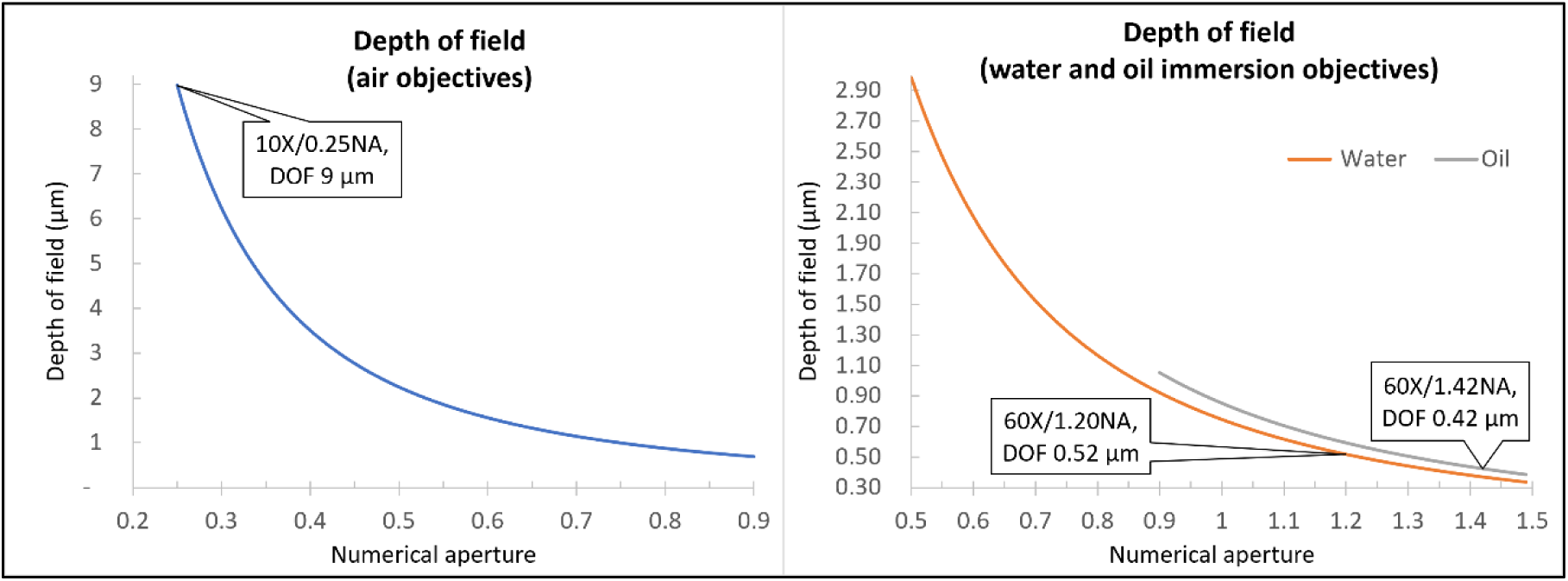
Depth of field vs numerical aperture. **a)** DOF vs NA of air objectives. The call-out sign illustrates the DOF of the 10X/0.25NA objective used in this study, which extends up to 9 µm. **b)** DOF vs NA of oil and water immersion objectives used in this study. The call-out texts show a DOF of 0.52 µm for the 60X/1.2NA water immersion objective and a DOF of 0.42 µm for the 60X/1.42NA oil immersion objective.

### S4. Penetration depth vs refractive index

In total internal reflection fluorescence (TIRF) microscopy, the sample is illuminated by a thin evanescent field originating at the interface between the sample media and the core material used as a sample substrate (Supplementary Figure S4a). The illumination intensity decays exponentially according to the formula *I*(*z*) = *I*_0_*e*^−*z*/*d*^, where *I*_0_ is the intensity at the sample-substrate interface, *z* is the distance in the z-direction perpendicular to the core material surface (sample substrate), and *d* is the depth of penetration of the evanescent field^44^. As a convention, the penetration depth of the evanescent field is defined as the distance at which the intensity equals 1/e (roughly, 37%) of the surface intensity (I_0_). The penetration depth (*d*) is given by the formula 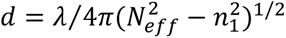, where λ is the excitation wavelength, *n*_1_ is the refractive index of the sample media, and *N*_eff_is the effective refractive index of the core material (sample substrate). For glass-based TIRF, this last term is given by *N*_*glass*_ = *n*_2_*sin*θ, where θ is the illumination angle, and *n*_2_ is the refractive index of the glass substrate. Thus, the penetration depth of the evanescent intensity for glass-based TIRF is given by the formula 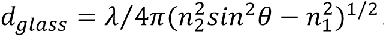. Similarly, for multimode waveguide-based TIRF, the penetration depth 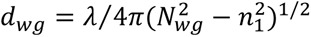 is dependent on the effective refractive index of the core material *N*_*wg*_, which varies for each guiding mode^82^. Typically, the *N*_*wg*_ values are solved by numerical simulations.

Supplementary Figure S4b provides a theoretical simulation of the penetration depth as a function of the effective refractive index used for TIRF microscopy. In this example, we considered a refractive index *n*_1_ ≈ 1.4 for the sample media, an excitation wavelength λ = 561 nm and, for glass-TIRF, a fixed illumination angle θ = 75 degrees. Therefore, for borosilicate glass (*n*_*glass*_ ≈ 1.52), an effective refractive index *N*_*glass*_ ≈ 1.47 was considered. Similarly, based on numerical simulations, for chip-TIRF, we assumed an effective refractive index *N*_*Si3N*4_ ≈ 1.75. In this configuration, glass-TIRF provides a theoretical penetration depth of ∼101 nm, while the high refractive index of the waveguide core material used in this study (Si_3_N_4_) allows for a theoretical penetration depth of ∼43 nm. By employing higher refractive index materials on the waveguide core, such as titanium dioxide (TiO_2_, *N*_*Tio2*_ ≈ 2.5), the penetration depth can be further narrowed down to ∼20 nm (ref. 83).

Although sub-100 nm penetration depth can be achieved on glass-based TIRF approaches via high illumination angles, these methods pose other limitations for clinical applications. In objective-based TIRF, for example, high-numerical aperture objectives (NA ≥ 1.4) are required for achieving the high illumination angles necessary for evanescent field excitation. As a consequence, high magnifications (≥ 60X) are also necessary, therefore restricting the exploitable TIRF field of view to around 50 µm × 50 µm. While prism-based TIRF allows fluorescence imaging with conventional objectives by decoupling the illumination and the collection paths, these approaches require precise optical alignment to control the penetration depth extent, contrary to the photonic chip, where ultrathin optical sectioning is achieved irrespective of the coupling precision.

**Supplementary Figure S4.**
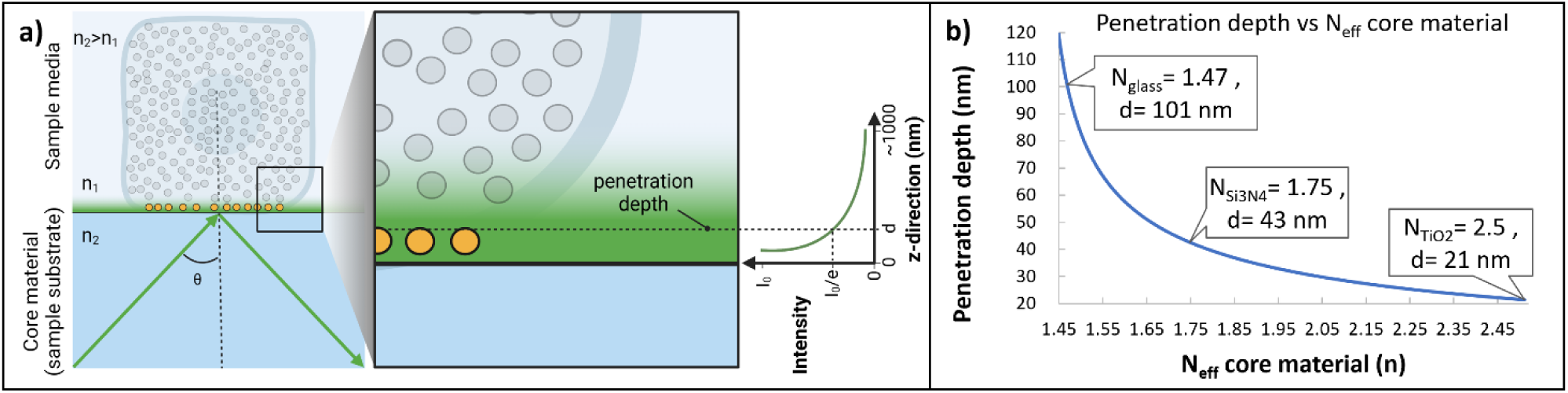
Penetration depth in TIRF microscopy. **a)** Schematic representation of total internal reflection fluorescence (TIRF). A light beam (green arrow) travels in a core material (sample substrate) with a refractive index n_2_ toward a sample media with a lower refractive index n_1_. Due to Snell’s law, upon hitting the substrate-sample interface on an angle θ greater than the critical angle θ_*c*_ = θθ*n*^−1^(*n*_1_⁄*n*_2_), the light beam is reflected into the core material medium. At the core material surface, however, a decaying field known as evanescent field appears, enabling the excitation of fluorescent molecules in its reach. The penetration depth of the evanescent field is defined as the distance in the z-direction perpendicular to the core material surface at which the illumination intensity equals 1/e of the illumination intensity (I_0_) at the sample-core interface. **b)** Theoretical simulation of the penetration depth as a function of the effective refractive indices of the core material. Here, the high index of refraction of the chip provides ultrathin optical sectioning as compared to glass-based TIRF systems.

### S5. Sample preparation workflow

Photonic chip-based microscopy is compatible with the conventional histology workflow of FFPE sections. Supplementary Figure S5 illustrates the sample preparation steps for chip-based microscopy of FFPE sections. The lateral dimensions (roughly, 2.5 cm × 2.5 cm) of the photonic chips used in this study made it necessary for manual handling of the sample throughout the complete preparation process. However, the photonic chip can be manufactured in identical sizes as conventional glass slides, making it possible for automated sample preparation via commercially-available tissue processing and immunostaining devices.

**Supplementary Figure S5.**
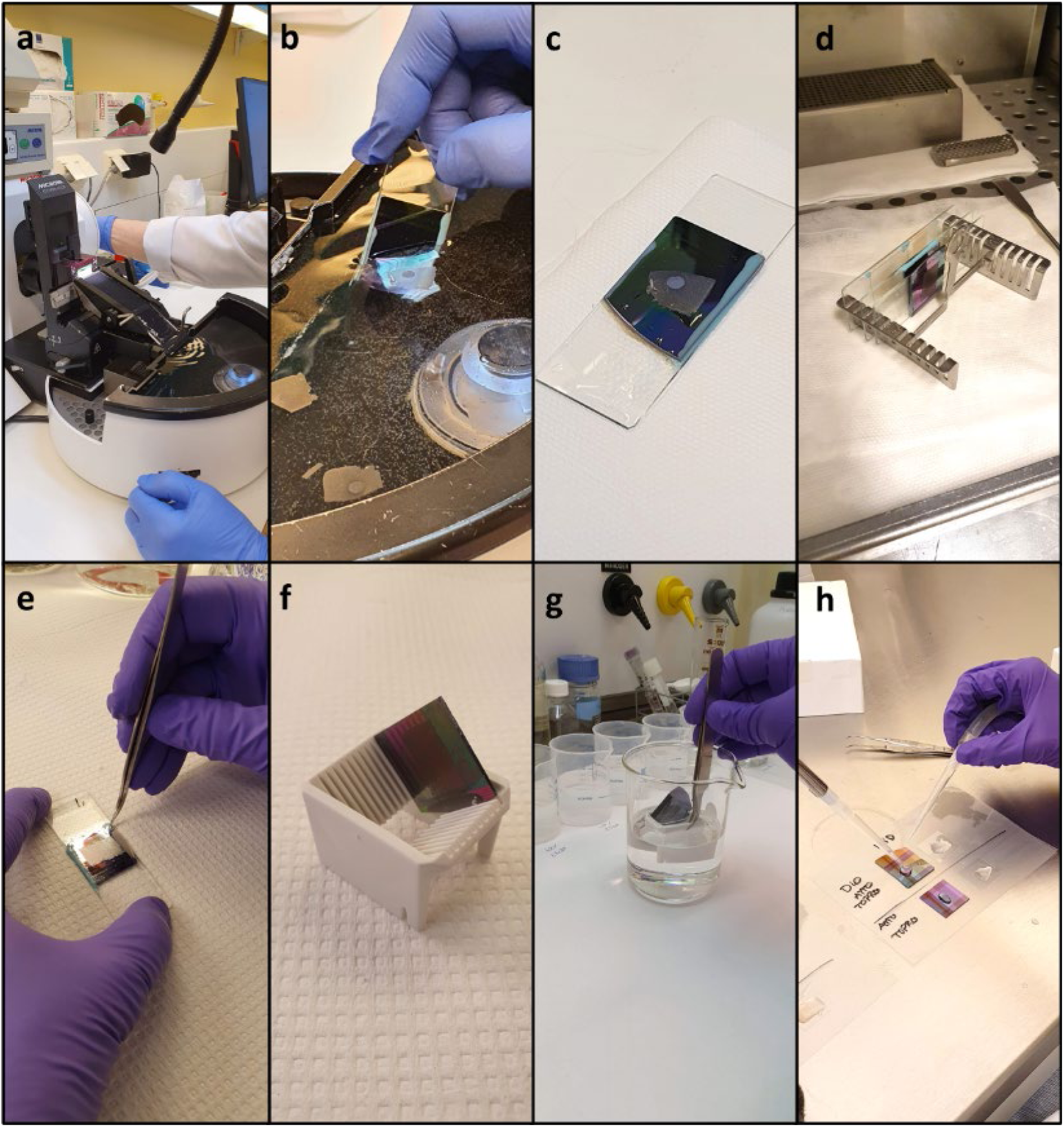
Sample preparation workflow for on-chip FFPE histology. **a)** The paraffin block is sectioned on a microtome equipped with a water bath. **b)** Upon sectioning, the sample is floated on the water bath and further scooped with a photonic chip previously glued to a conventional glass slide. See details in Supplementary Video V3. **c)** The photonic chip is placed on a flat surface at room temperature for an hour, for drying. **d)** The photonic chip is placed inside a 60 °C oven for overnight melting of the paraffin. **e)** The photonic chip is carefully detached from the glass slide by peeling off the picodent twinsil glue. **f)** The photonic chip is placed on a coverslip rack. **g)** The photonic chip is immersed in several reagents for deparaffinization and rehydration of the sample. **h)** The sample is fluorescently labeled using a single-channel micropipette and an aspirator pump.

### S6. Mouse kidney immunolabeling

The photonic chip is compatible with immunolabeling approaches. In this example, a zinc-fixed paraffin-embedded (ZnFPE) mouse kidney section was fluorescently immunolabeled using recombinant anti-alpha smooth muscle actin (⍺;-SMA) antibody (ab124964, Abcam). To enable optimal immunolabeling, after deparaffinization and rehydration (see Supplementary Information S5), the photonic chip was immersed in citrate buffer and microwave-heated for ten minutes at maximum power. For a contextual visualization, the sample was also labeled against membranes. Supplementary Figure S6 shows the ZnFPE mouse kidney sample in different levels of magnification (10X and 20X). The ⍺;-SMA is displayed in cyan and the membranes are shown in magenta.

**Supplementary Figure S6.**
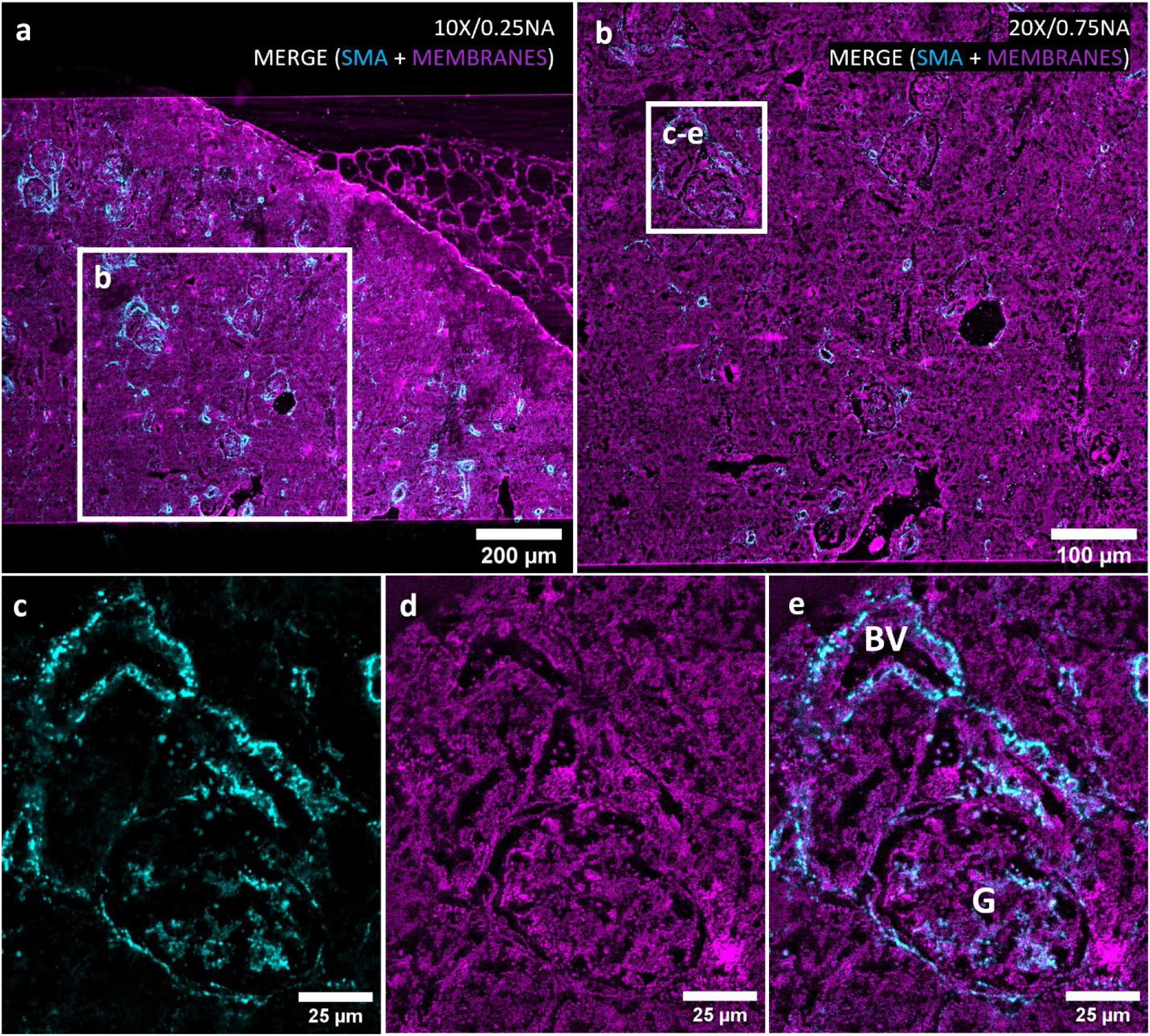
Photonic chip-based immunolabeling of paraffin sections. **a)** 10X/0.25NA chip-TIRF image of a ZnFPE mouse kidney showing ⍺;-SMA in cyan and membranes in magenta. The white box denotes the region further described in b). **b)** 20x/0.75NA chip-TIRF image of the mouse kidney section. The white box denotes a sample region containing a glomerulus and an artery. **c-e)** A zoom-in view of the selected region allows for detailed visualization of the sample structure. BV: blood vessel; G: glomerulus.

### S7. Magnified view of the H&E colorectal sample

Magnified view of the adenocarcinoma region (Ad) in the colorectal sample.

**Supplementary Figure S7.**
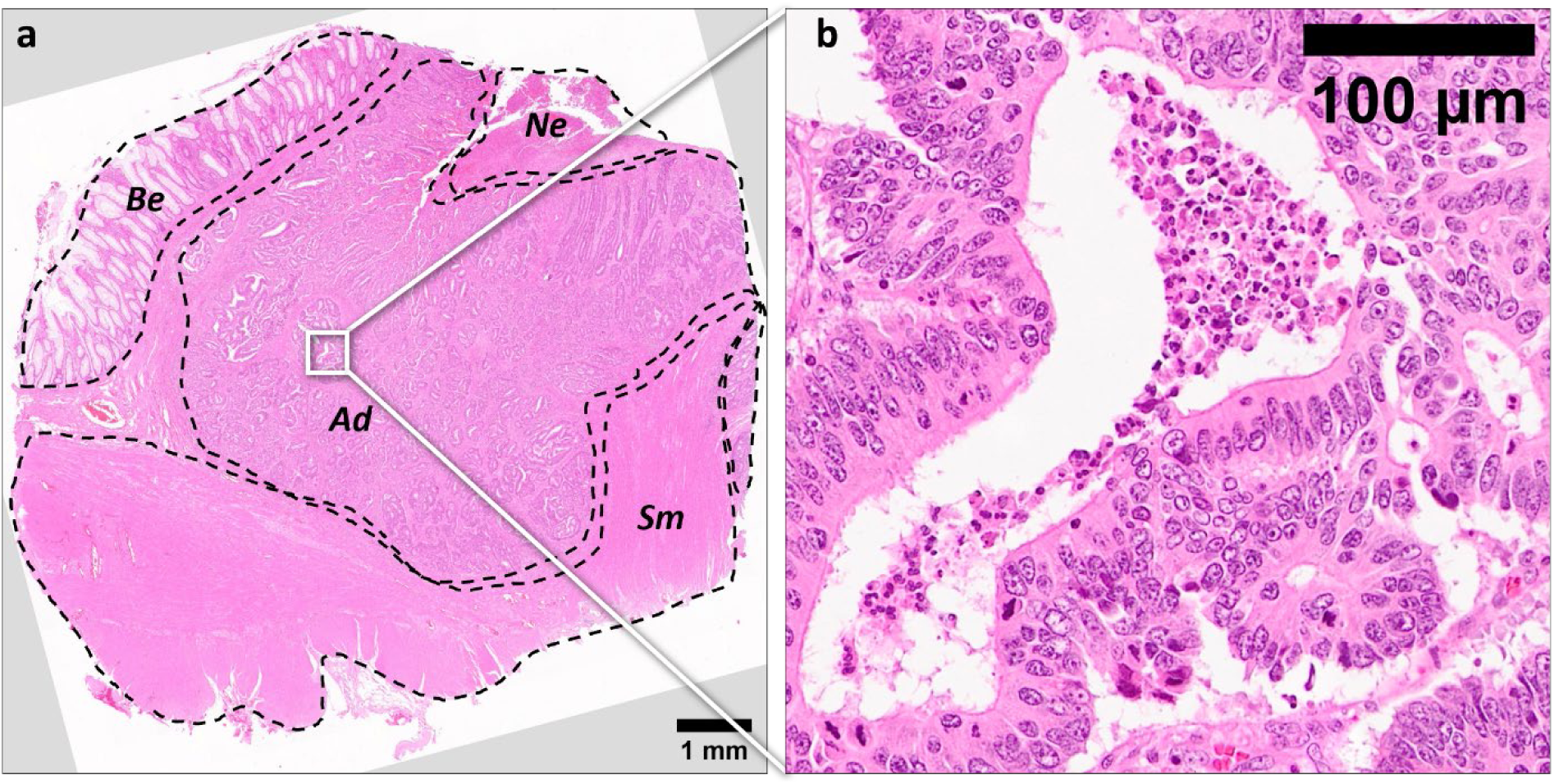
Zoom-in view of the adenocarcinoma region (Ad). The high magnification allows the distinction of structural features (glands lined with pseudostratified cancer cells showing varying degrees of pleomorphism and a lumen with necrotic debris) in the colorectal sample that are not distinguishable in low magnification (Figure 3a).

### S8. Multicolor chip-TIRF image acquisition

Supplementary Figure S8 illustrates the acquisition steps for a multicolor chip-TIRF image of an FFPE prostate tissue section using a 10X/0.25NA collection objective. After selecting the region of interest, consecutive waveguides are imaged in chip-TIRF modality (Supplementary Figure S8a). Thereafter, the membrane images are stacked and projected using the maximum projection option in FIJI. The process is repeated for a different labeled channel, in this case, nuclei (Supplementary Figure S8b), using a specific excitation laser wavelength and corresponding emission filter. Finally, the membrane and nuclei images are merged in FIJI to obtain a multicolor chip-TIRF image.

**Supplementary Figure S8.**
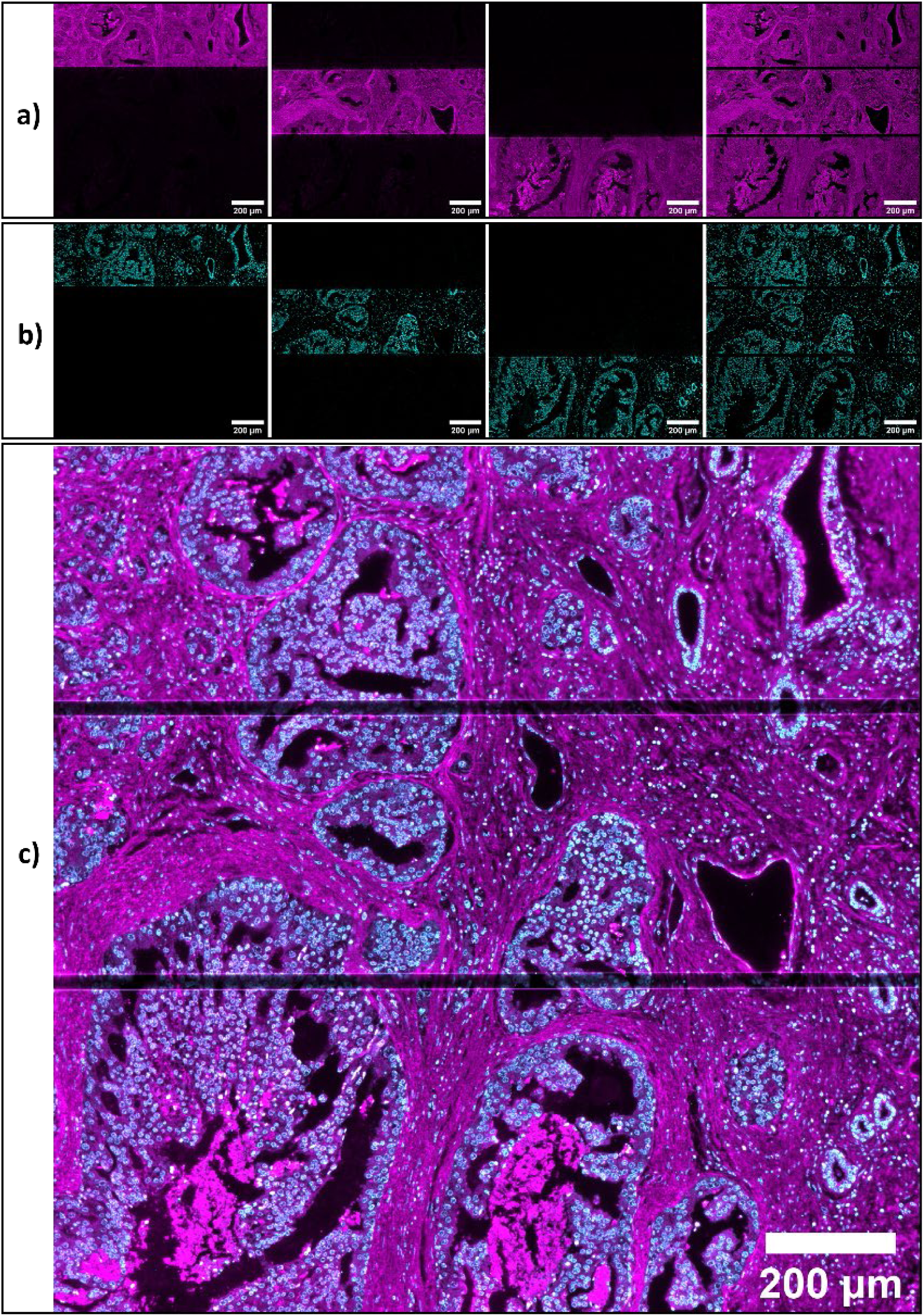
Multicolor chip-TIRF acquisition. **a)** Three individual images of membranes are collected in chip-TIRF modality and further projected into a single image in FIJI. **b)** The process is repeated for the nuclei signal. **c)** Finally, the projected images are merged in FIJI to obtain a multicolor chip-TIRF image.

### S9. Decorrelation analysis 10X/0.25 images

A decorrelation analysis^64^ was used to quantify the resolution enhancement of chip-MUSICAL over conventional EPI fluorescence images (Supplementary Figure S9). Both chip-MUSICAL and EPI images were acquired with a 10X/0.25NA objective over the same field of view. By analyzing the images under a series of high-pass filters, the decorrelation algorithm computes the maximum spatial frequency at which it is still possible to distinguish between signal and noise, enabling resolution estimation of microscopy images. For this analysis, the MATLAB version of the algorithm was used. In both cases (i.e. EPI and chip-TIRF), the number of sample points (Nr) was set to 100 and the number of filters (Ng) to 50. The results revealed an estimated resolution of 1563 nm and 1310 nm for the EPI and chip-MUSICAL images, respectively, suggesting nearly a 1.2-fold resolution improvement of chip-MUSICAL over EPI. We acknowledge that the chip-based microscopy setup used in this study does not perform at the theoretical limit of diffraction due to minor imperfections along the optical path length of the system. Supplementary Figure S9 shows the corresponding plots for resolution estimation on each decorrelation analysis. The resolution is calculated with the formula 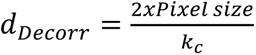, where *k*_*c*_ corresponds to the maximum normalized spatial frequency shown in the plot (e.g. *k*_*c*_ = 0.8316 for EPI, and *k*_*c*_ = 0.1984 for chip-MUSICAL).

**Supplementary Figure S9.**
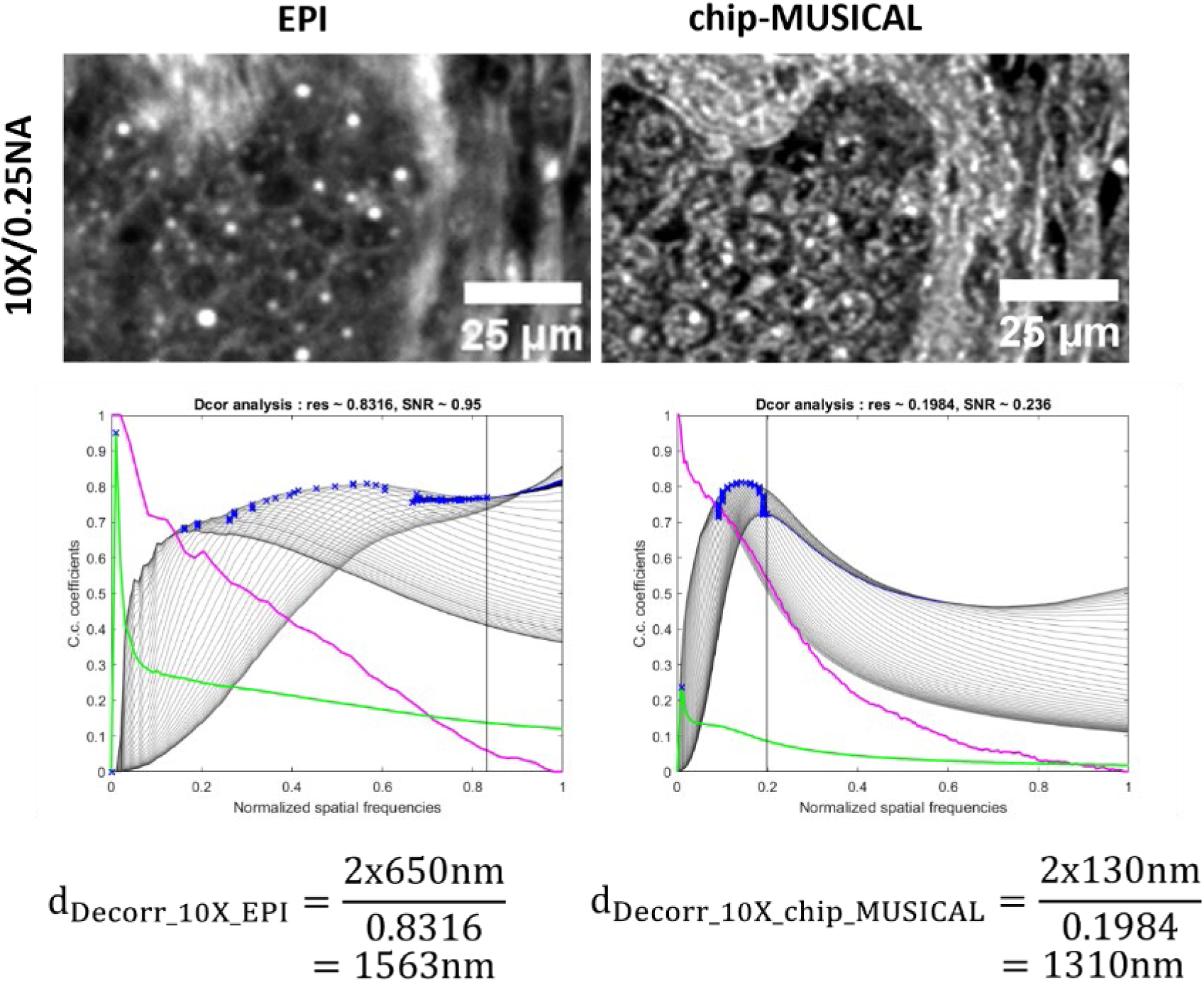
Decorrelation analysis of 10X/0.25NA EPI and chip-MUSICAL images. Photonic chip-based microscopy enables resolution improvement over conventional EPI fluorescence methods. In this example, the membrane channels of the FFPE prostate sample shown in Figure 4c and Figure 4d were used for resolution estimation via decorrelation method. The results obtained here revealed a resolution of 1563 nm and 1310 nm in EPI and chip-MUSICAL, respectively, which suggest a nearly 1.2-fold resolution improvement via chip-MUSICAL over EPI.

### S10. Decorrelation analysis 60X/1.2 images

Following the steps described in Supplementary Information S9, a decorrelation analysis was performed on the 60X/1.2NA EPI and chip-MUSICAL images corresponding to the membrane channels of Figure 4e and Figure 4f, respectively. Here, the decorrelation algorithm estimated a resolution of 461 nm and 194 nm for the EPI and chip-MUSICAL images, respectively, which suggests nearly a 2.4-fold resolution improvement of chip-MUSICAL over EPI. Supplementary Figure S10 shows the corresponding plots for resolution estimation on each decorrelation analysis. The resolution is calculated with the formula 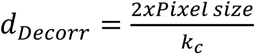, where *k*_*c*_ corresponds to the maximum normalized spatial frequency shown in the plot (e.g. *k*_*c*_ = 0.4677 for EPI, and *k*_*c*_ = 0.2222 for chip-MUSICAL).

**Supplementary Figure S10.**
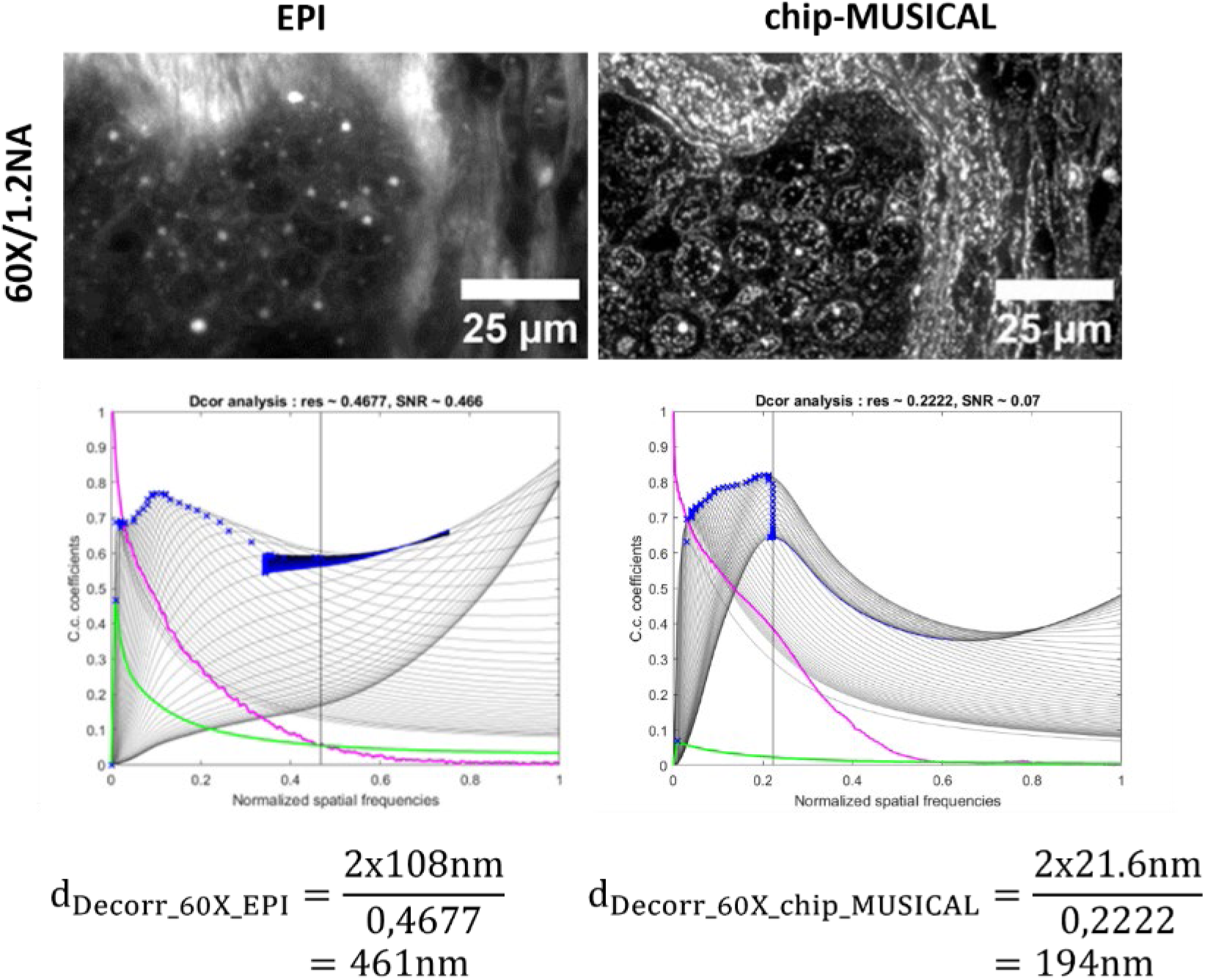
Decorrelation analysis of 10X/0.25NA EPI and chip-MUSICAL images. Photonic chip-based microscopy enables resolution improvement over conventional EPI fluorescence methods. In this example, the membrane channels of the FFPE prostate sample shown in Figure 4e and Figure 4f were used for resolution estimation via decorrelation method. The results obtained here revealed a resolution of 461 nm and 194 nm in EPI and chip-MUSICAL, respectively, which suggest a nearly 2.4-fold resolution improvement via chip-MUSICAL over EPI.

### S11. Advantages of photonic chip-based microscopy for FFPE histology

Photonic chip-based microscopy offers several advantages for histological analysis of FFPE sections. These include:

a. **Compatibility with standard histological workflows**. The photonic chip withstands all the steps necessary for sample preparation of FFPE sections, including water immersion for sample scooping, oven incubation at 60°C for paraffin melting, xylene immersion for deparaffinization, descendent alcohol incubations for sample rehydration, and heat-induced antigen retrieval for immunolabeling. In addition, the photonic chip can be fabricated to match the dimensions of standard microscopy glass slides, facilitating seamless integration with automated processing equipment used in clinical settings.
b. **High contrast over large fields of view**. The evanescent field provided by the photonic chip enables ultra-thin optical sectioning (<50 nm) of the FFPE samples along the entire length of the waveguide, supporting illumination on the centimeter scale. In addition, the decoupled excitation and collection light paths enable the acquisition of TIRF images using any arbitrary magnification objective lens, thus opening avenues for high-contrast images over large areas^21^, thus overcoming the field of view limitations of conventional TIRF systems, typically restricted to approx. 50 µm × 50 µm.
c. **Multimodal imaging**. The photonic chip enables diverse imaging modalities such as TIRF, FF-SRM, SMLM, SIM, and CLEM^25, 27, 29, 56, 71^. Also, the chip fabrication supports the inclusion of landmarks that can further aid in the identification of specific regions of interest across different microscopy methods including atomic force microscopes (AFM) and scanning electron microscopes (SEM), as illustrated in Figure 5. In addition, the photonic chip is compatible with complementary imaging tools such as microfluidics^74^, which are often used in multi-omic research.
d. **Support of high-spatial frequency illumination**. The high refractive index of the waveguide core material (*n* ≈ 2 for Si_3_N_4_) supports much higher spatial frequencies than supported by conventional free-space illumination, which further assists in an improved lateral resolution of FF-SRM methods^25^. While in EPI illumination approaches the highest spatial frequency illuminating the sample is limited by diffraction according to the excitation wavelength and the numerical aperture of the objective lens, the spatial frequencies supported by the waveguide material on the photonic chip platform are determined by the effective refractive index of the waveguide material, allowing for high-spatial frequency illumination beneficial for SRM^25^.
e. **Compact footprint and retrofittable**. Photonic chip-based microscopy can be integrated into standard upright microscopes upon a few adaptations, making it an attractive solution for super-resolution multimodal imaging of tissue sections. In addition, the system operability allows for a quick and simple adoption for non-expert users of super-resolution fluorescence-based microscopy methods. Also, the photonic chips can be mass fabricated following standard semiconductor techniques and re-utilized in multiple assays, which makes them a suitable option for routine laboratories.

### S12. FFPE placental sample detachment

An indispensable requisite for successful TIRF imaging is sample adhesion. Throughout this study, we faced sample micro-detachment issues that hampered the imaging capabilities of the chip-TIRF technique. When these micro-detachments occurred, discontinuities in the chip-TIRF signal were observed. Supplementary Figure S12 illustrates a sample micro-detachment of an FFPE placental section. The white-dotted oval in Supplementary Figure S12a denotes a region of a chip-TIRF image with signal discontinuity. Further observation on a scanning electron microscope (Supplementary Figure S12b) reveals a sample micro-detachment at the same location.

**Supplementary Figure S12.**
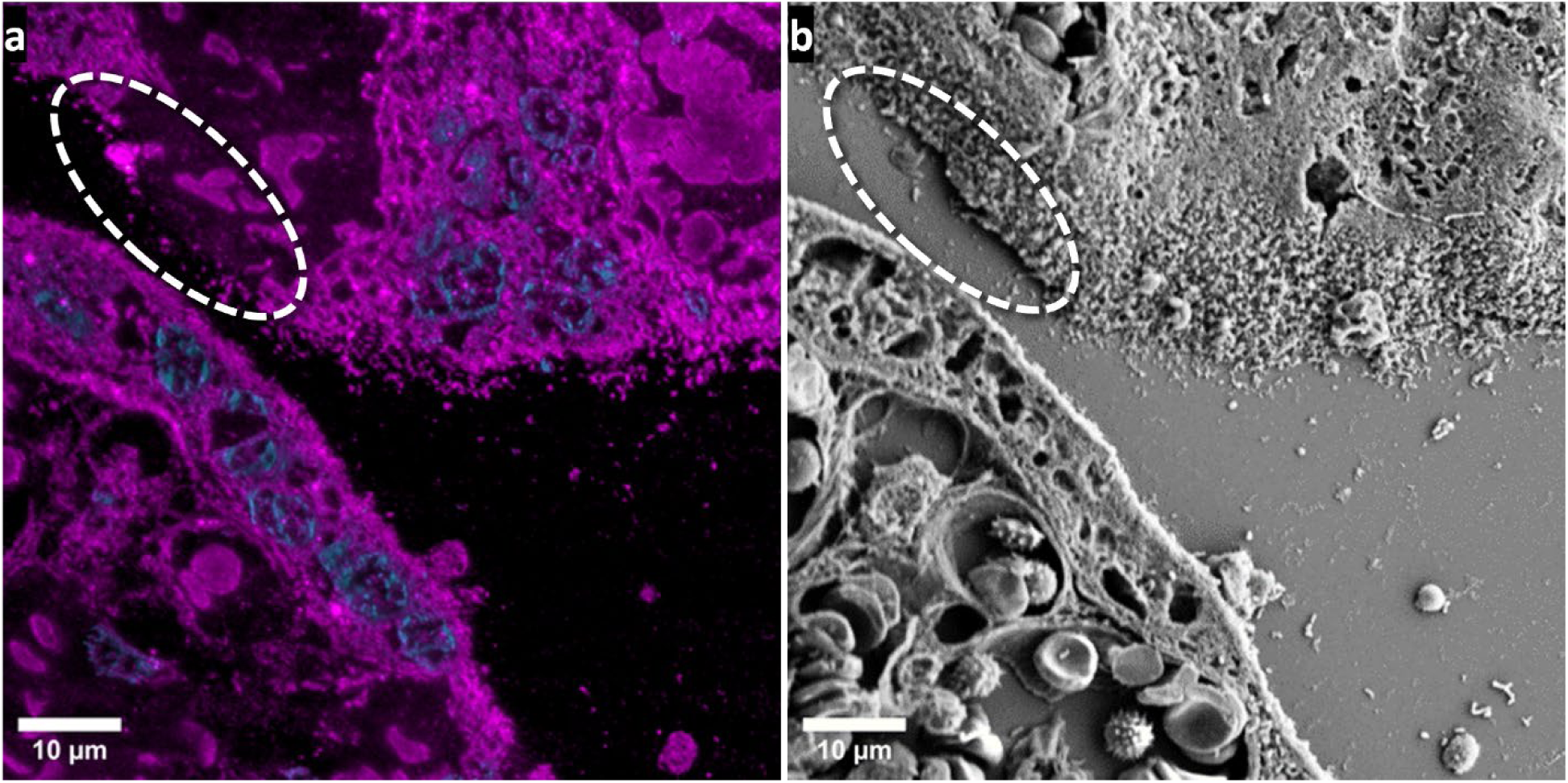
Detachment example of a placental FFPE sample on a photonic chip. **a)** The white-dotted oval denotes a region of a chip-TIRF image with signal discontinuity. **b)** Further observation under a scanning electron microscope (SEM) revealed a sample micro-detachment at the same region.

### S13. Optimization steps for on-chip FFPE tissue imaging

The main experimental challenge of this study was sample detachment. While photonic chip-based microscopy enables superior optical sectioning, it requires the tissue section to lay perfectly flat on the waveguide surface to enable fluorescent emission via evanescent field excitation. As a consequence, micro-detachment gaps greater than the evanescent field depth render dark patches along the chip-TIRF image that hamper the visualization of the samples (see Supplementary Information S12). We tried to address this issue in different ways including a) mechanical pressure, b) limited sample hydration, and c) chip coating. In the first attempt, mechanical pressure, we aimed at pushing down the detached FFPE sample to establish contact with the waveguide surface. To achieve this, we followed the chip-based fluorescence labeling process as outlined in the Materials and Methods section, and then placed the chip in between a ferromagnetic plate at the bottom and four cylindrical magnets of 1 mm in diameter on the corners of the coverslip to exert uniform pressure over the tissue section. This method, however, did not improve the detachment issues of the FFPE samples. To further understand the detachment issue, we performed a topographical assessment of deparaffinized tissue samples using atomic force microscopy (AFM). In this process, we found out that FFPE tissues exhibited swelling along with the rehydration process (see Supplementary Information S14). This brought us to our second attempt, limiting the sample hydration to a maximum ratio of 5:95 H_2_O to ethanol, to minimize the chances of swelling and, therefore, reduce the sample detachment. Despite the effort, we observed no significant differences in sample detachment compared to further hydrated samples in 30:70 H_2_O to ethanol. A plausible explanation for this is that, despite the limited H_2_O content in the rehydration steps, the samples inevitably underwent further incubation steps in aqueous media for the fluorescence labeling and washing, thus absorbing more H_2_O and, subsequently, swelling. Finally, we explored functionalizing the chip surface to ensure sample adhesion through the preparation process. Here, we tested various coating alternatives including poly-L-lysine, histogrip, and protected isocyanate^84^ (see Supplementary Information S15). While we observed a better sample adhesion compared to uncoated chips, we could not appreciate significant differences between the coating strategies. Instead, we noticed that the detachment issues were more dependent on the sample type than on the coating method. For example, in the colorectal sample, the adenocarcinoma area (Figure 3) showed optimal adhesion while the benign epithelium region showed significant detachment (see Supplementary Information S14). Similarly, the placental section exhibited more detachment issues along the apical side of the chorionic villi, in comparison with the stromal region (see Supplementary Information S11 and S15). Hence, we opted for the easiest and most affordable coating option, poly-L-lysine, to carry out this study.

The second challenge we encountered was labeling. In the attempt to find a membrane marker that could give us a contextual visualization of the sample, we tried the different variations of the CellMask dye family (CellMask Green, CellMask Orange, and CellMask Deep Red), as well as the Vybrant DiI and DiO fluorophores, but all these showed high affinity to the waveguide surface, which led to a high background signal during imaging acquisition^30^. We tried to address this issue by using blocking reagents such as bovine serum albumin and increasing the number of washing steps after fluorescence labeling, but we observed no reduction in the background signal. Interestingly, we found out that the MitoTracker Deep Red FM (MTDR) rendered unspecific labeling of paraffin-embedded samples while producing negligible background on the waveguide surface. We acknowledge that the MTDR is intended for live cell analysis of mitochondria. However, the results obtained in this study suggest that this marker serves the purpose of contextual visualization as eosin does in the H&E staining. For example, the comparative view of the colorectal sample showed a clear correspondence between the pink areas in Figure 3a, and the magenta structures in Figure 3b. In addition, the CLEM imaging of the placenta (Figure 5) confirmed that the labeled structures in magenta color matched the features observed in the SEM image.

Lastly, we faced issues with unguided light. In the initial imaging attempts, we experienced unwanted side illumination along the sample volume stemming from uncoupled light traveling in free space and reaching the sample region. This issue severely affected the contrast of the chip-TIRF images, hindering the imaging capabilities of the system. We successfully blocked the unguided light by incorporating a small piece (∼1 × 1 × 10 mm) of custom-made black polydimethylsiloxane (PDMS) near the coupling facet of the photonic chip, which served as shielding against the unguided light.

### S14. Atomic force microscopy (AFM) analysis of FFPE sections

In this study, we observed a heterogeneous behavior of the micro-detachment phenomena (see Supplementary Information S12). For example, in the FFPE colorectal sample (Supplementary Figure S14a), the benign colonic epithelium region became commonly micro-detached after sample preparation. While the sample was still optimal for EPI fluorescence microscopy (Supplementary Figure S14b), the signal discontinuities on the chip-TIRF image (white arrows in Supplementary Figure S14c) suggested signal micro-detachments of the cell walls of the enterocytes in this region. To further understand the root cause of such micro-detachments, we performed a height characterization over a colonic crypt section of an FFPE colorectal tissue using an atomic force microscope (AFM) (Supplementary Figure S14d). For this, we set the AFM to quantitative imaging mode and measured the height of the sample under 95 % ethanol incubation to enable a topographical visualization of the sample in a dehydrated state (Supplementary Figure S14e). Thereafter, we repeated the measurement over the same region, this time under 50 % ethanol incubation, to visualize the morphological changes of the sample induced by hydration (Supplementary Figure S14f). A line profile measurement over the same region revealed significant morphological changes on the sample, particularly over the lumen of the enterocytes, suggesting that this vacuolated region undergoes swelling upon hydration (Supplementary Figure S14g). We also performed force measurements on the colorectal sample (not shown) and found that the cell wall of the enterocytes was significantly stiffer than the vacuolated regions exhibiting swell. Although the root source of the sample micro-detachment phenomena remains unclear, we hypothesize that, upon hydration, the vacuolated structures undergo swelling, expanding in all directions and therefore pulling out the enterocytes’ cell wall from the chip surface.

**Supplementary Figure S14.**
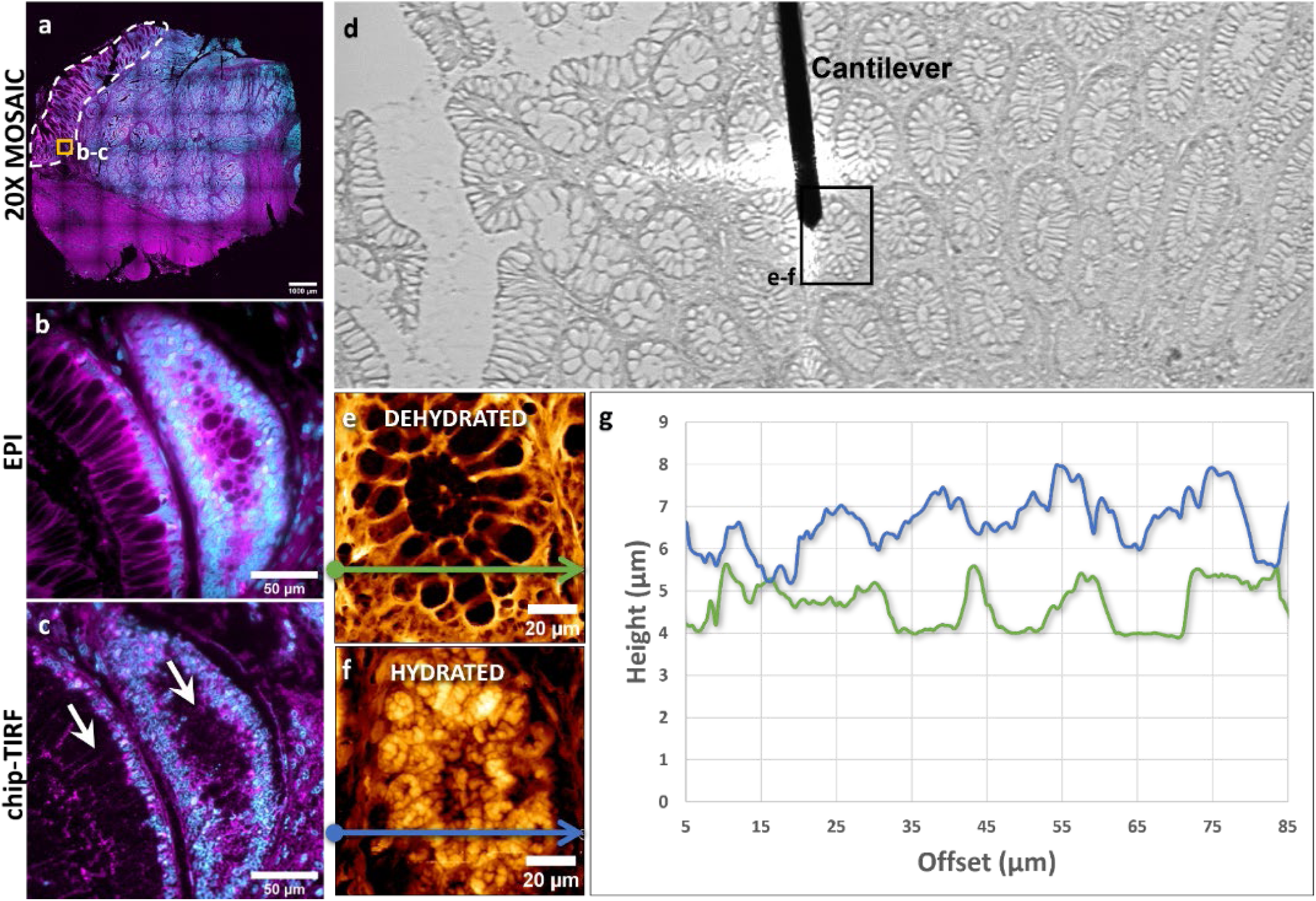
AFM measurements on an FFPE colorectal tissue section over different hydration stages. **a)** EPI fluorescent mosaic image of a colorectal sample. The white-dotted line denotes the benign colonic epithelium region. The yellow box denotes the regions imaged in b) and c), respectively. **b)** EPI fluorescent image of a colonic epithelium. Membranes are displayed in magenta and nuclei in cyan. **c)** chip-TIRF image of the same region. The white arrows illustrate the regions with signal discontinuity due to sample micro-detachment. **d)** Wide-field image of a colorectal sample on an AFM system. The black box denotes the region of the sample measured in e) and f) under different hydration conditions. **e)** Height measurement of a colonic crypt under 95 % ethanol incubation. The green line denotes the direction of the line profile measurement performed over the dehydrated sample. **f)** Height measurement of a colonic crypt under 50 % ethanol incubation. The blue line denotes the direction of the line profile measurement performed over the hydrated sample. **g)** The line profile measurements show significant swelling of the FFPE sample in a hydrated state as compared to the dehydrated state.

### S15. Coating strategies for on-chip FFPE sample adhesion

To improve the FFPE sample adhesion to the photonic chip, diverse coating methods were tested, including histogrip (cat. # 8050, ThermoFisher Scientific), protected isocyanate^84^ and poly-L-lysine coating (cat. # P8920, Sigma-Aldrich). In this case, three chips were used for the comparison, each one with a different coating. An individual FFPE placental section from the same block was placed on top of each coated chip and further labeled as described in the Materials and Methods section. To enable a comparative view, the samples were imaged both in EPI (top view) and chip-TIRF (bottom view). Supplementary Figure S15 illustrates the adhesion effect of different coatings on photonic chips. While all coating agents successfully retained the samples on the chip, we observed sample micro detachments that hampered the imaging capabilities of the photonic chip (yellow arrows on the bottom panels). Having found no significant differences among the three coating methods, in this work, we opted for the most user-friendly and economical method of all three, namely, the poly-L-lysine coating.

**Supplementary Figure S15.**
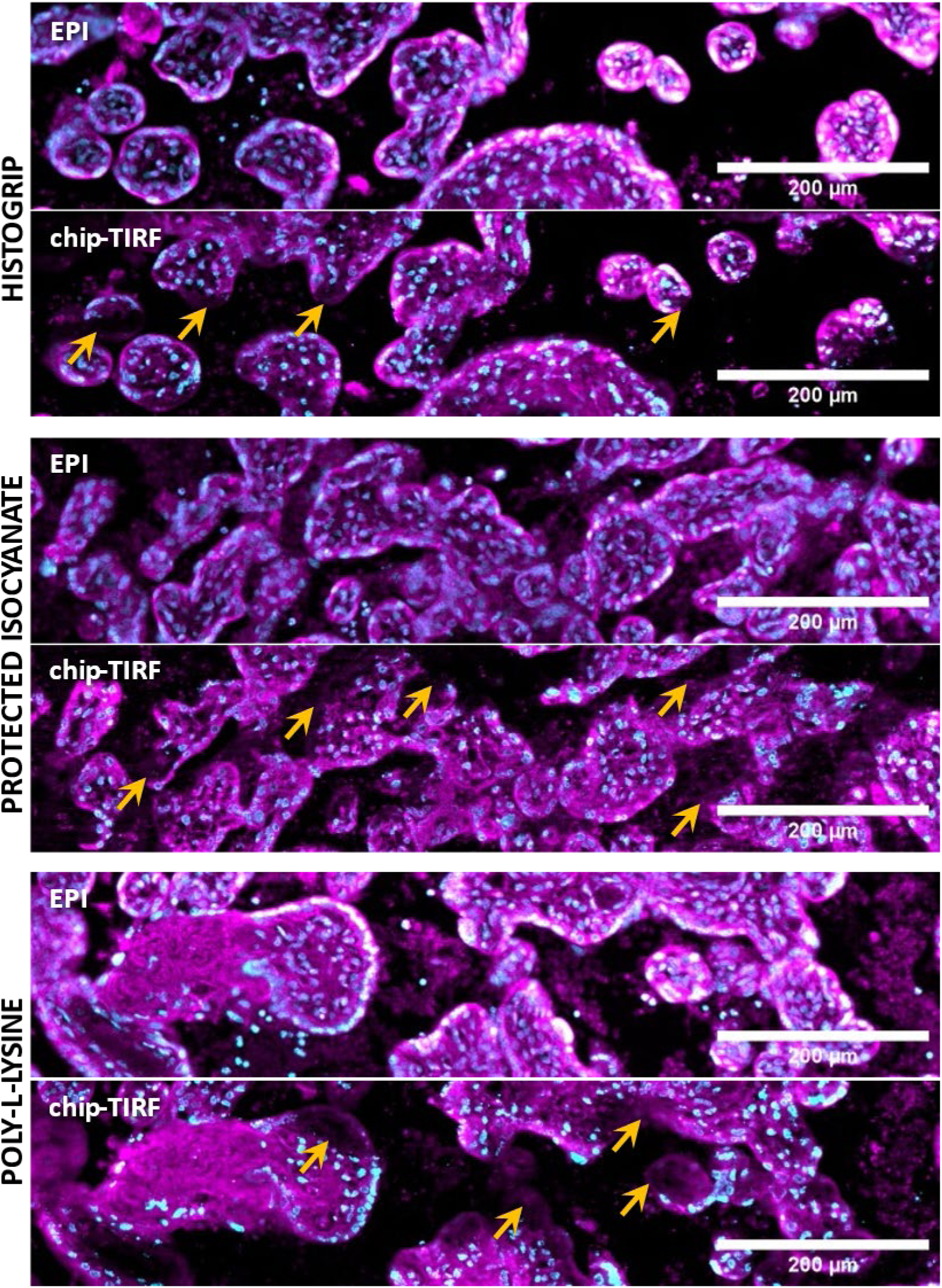
Coating strategies for on-chip FFPE sample adhesion. Despite retaining the FFPE samples on the chip, none of the coating methods tested, namely, histogrip, protected isocyanate (PI), and poly-L-lysine (PLL) provided full sample adhesion. Instead, sample micro-detachments were observed in all of them.

### S16. Materials and reagents used for on-chip FFPE fluorescence labeling

Supplementary Table S16 provides a detailed description of the materials and reagents used for on-chip FFPE sample preparation.

**Supplementary Table S16.** Materials and reagents used for on-chip FFPE sample preparation.

| Material/<br>reagent | Manufacturer | Catalog<br>number | Stock<br>concentration | Working<br>concentration | Purpose |
| --- | --- | --- | --- | --- | --- |
| #1.5 coverslip | VWR | 631-0125 | - | - | 22 mm x 22 mm #1.5 coverslip |
| Picodent twinsil | Picodent | 1300 1000 | - | 1:1 mixture of<br>solution A and B | Dental cement. Gluing and<br>sealing. |
| Poly-L-lysine | Sigma-Aldrich | P8920 | 0.1 % (w v<sup>-1</sup>) in<br>H<sub>2</sub>O | 1:1 | Chip-surface coating for<br>improved adhesion of<br>biological samples |
| MitoTracker™ Deep<br>Red FM | Invitrogen | M22426 | 1 mM | 1:2000 | Unspecific staining for<br>contextual information<br>(similar to membrane dyes) |
| Sytox Green | Invitrogen | S7020 | 5 mM | 1:1000 | Nuclear staining |
| Phosphate-buffered<br>saline (PBS) | Sigma-Aldrich | D8662 | - | 1:1 | Washing steps |
| Glycerol | Sigma-Aldrich | G5516 | - | 1:1 | Mountant medium |
| Xylenes | Sigma-Aldrich | 214736 | - | 1:1 | Sample deparaffinization |
| Ethanol absolute | VWR Chemicals | 20821.296 | >99.8 % | 1:1 | Sample rehydration |
| Ethanol 96 % | VWR Chemicals | 20823.362 | 96 % | 1:1 | Sample rehydration |

### V1. chip-TIRF mode-averaging

Link: https://uitno.box.com/s/ycaj9q50q6526bls4ah7f4ab91ovy30o

**Supplementary Video V1.**
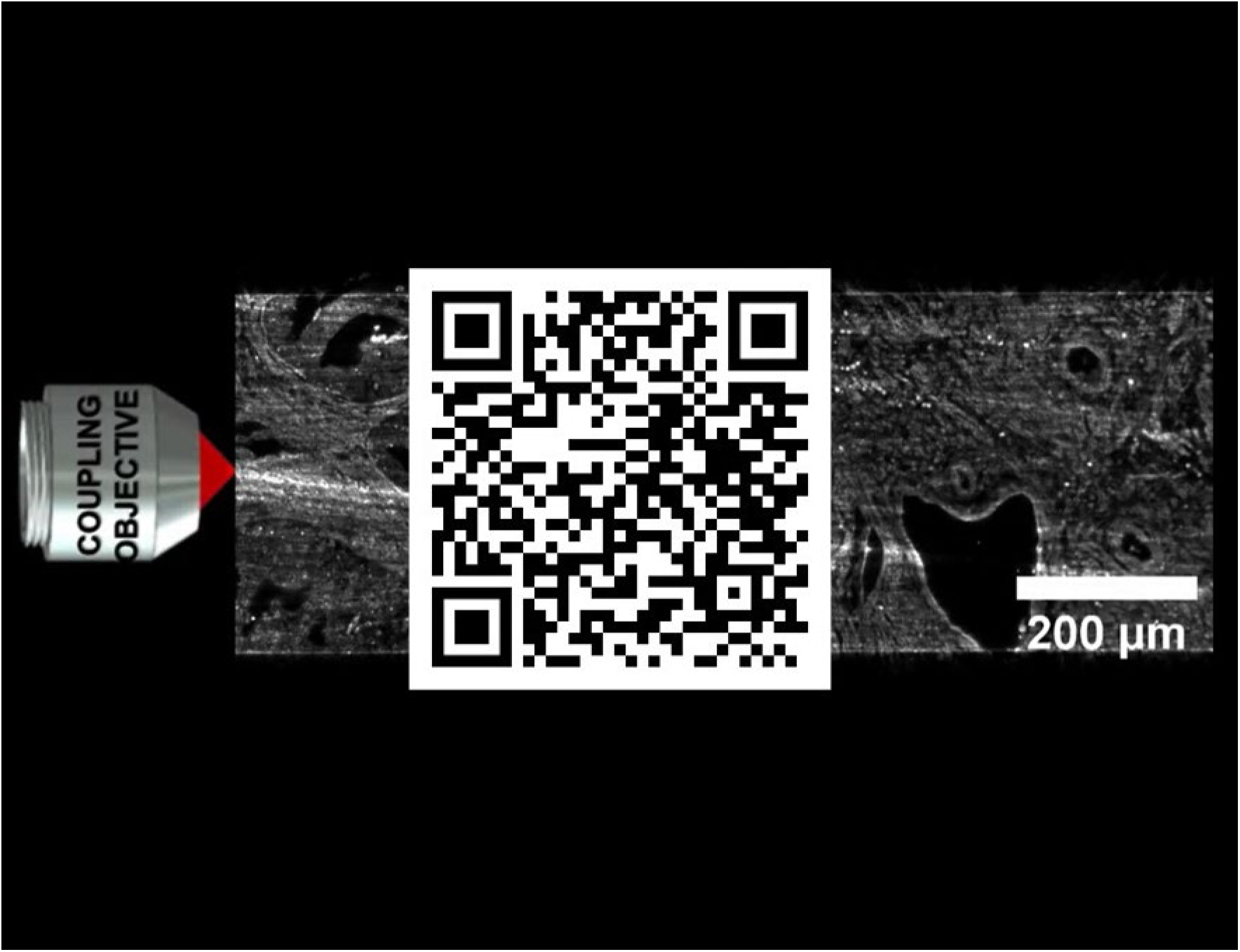
Chip-TIRF mode-averaging.

### V2. Comparison between EPI-MUSICAL and chip-MUSICAL

Link: https://uitno.box.com/s/rxsne0m3pdtgl3n3caj0rc6bvw4rqa0e

**Supplementary Video V2.**
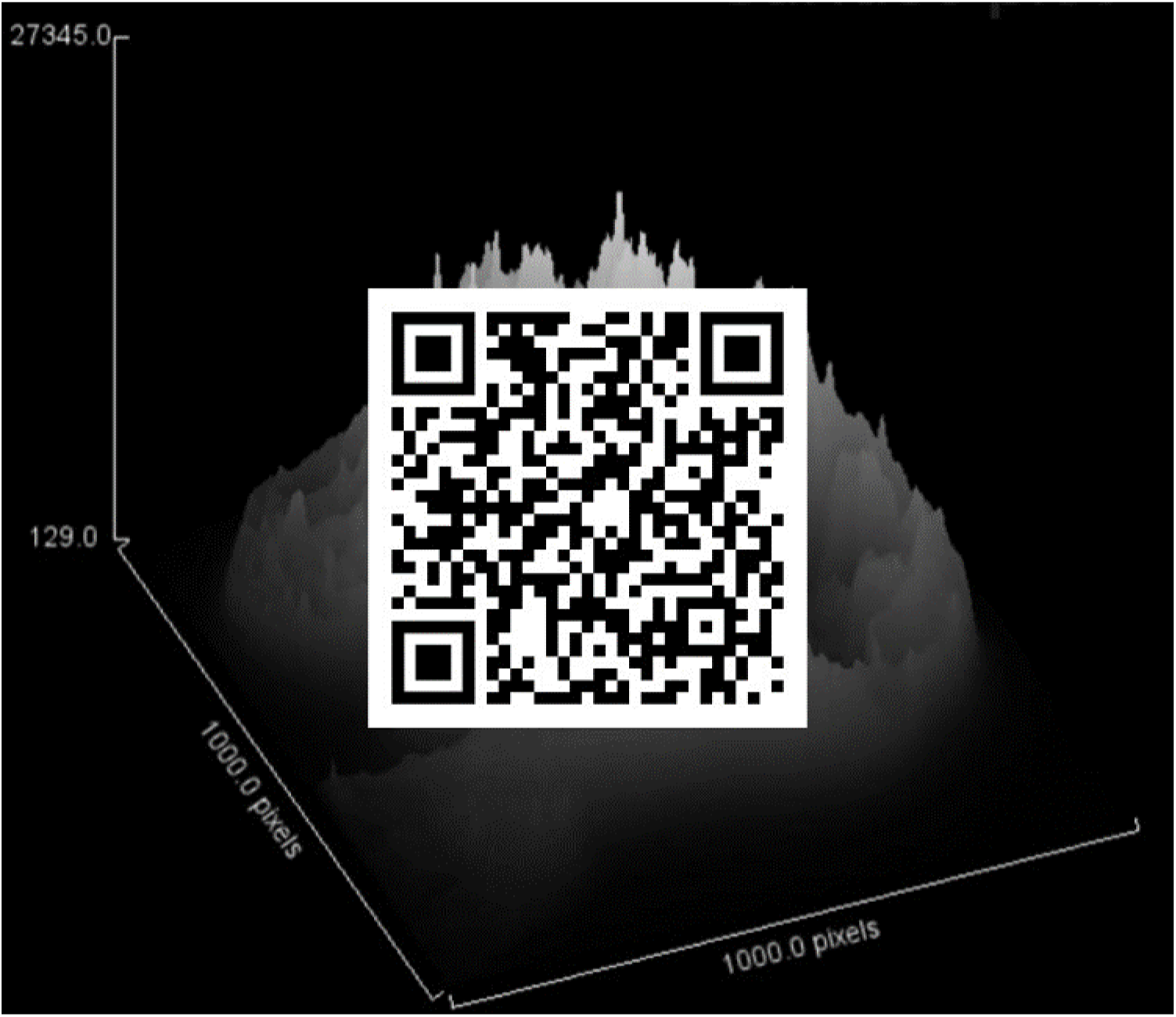
EPI-MUSICAL vs chip-MUSICAL comparison.

### V3. On-chip tissue scooping

Link: https://uitno.box.com/s/5cbrbpd1vyus4i3dmojgw89afdnvxafd

**Supplementary Video V3.**
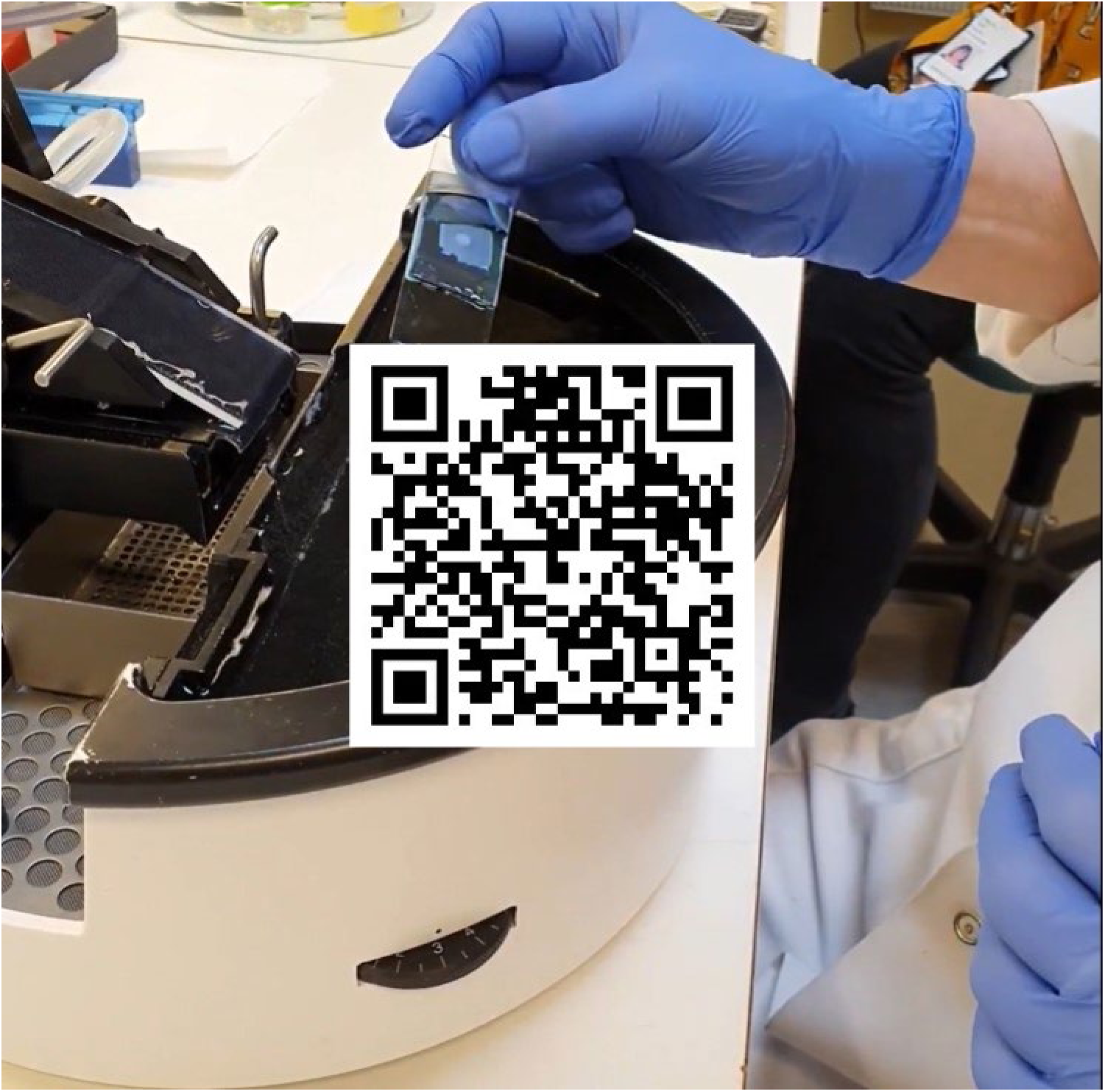
On-chip FFPE tissue scooping.

### V4. EPI vs chip-TIRF over large FOV

Link: https://uitno.box.com/s/vr3d0h4ku5y5q2irn4at5vgrlxgq4c2u

**Supplementary Video V4.**
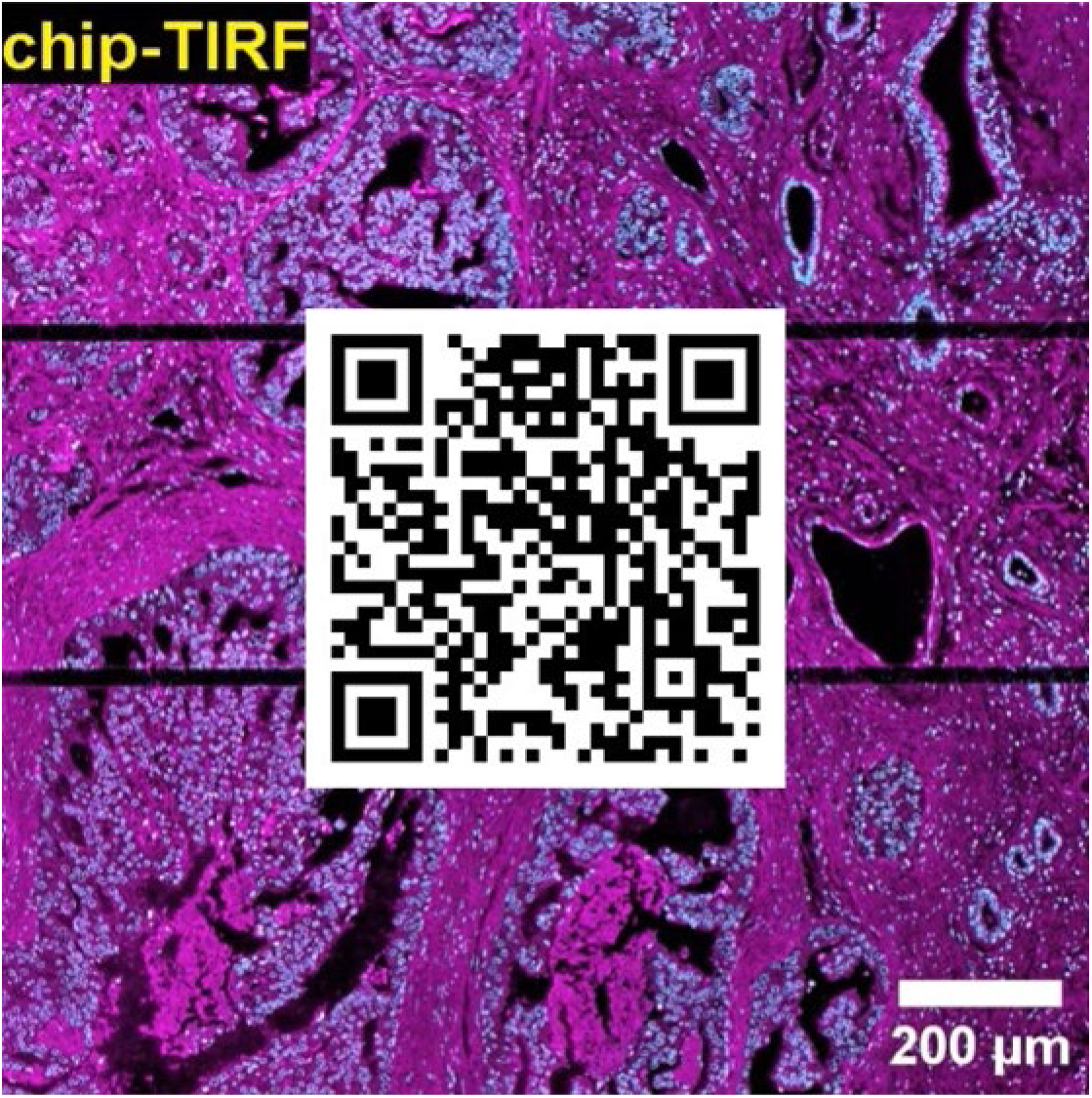
EPI vs chip-TIRF over large FOV.

### V5. FFPE CLEM imaging over large fields of view

Link: https://uitno.box.com/s/9kl4x0lq7gqst6d1vnr5nrqvyebfofef

**Supplementary Video V5.**
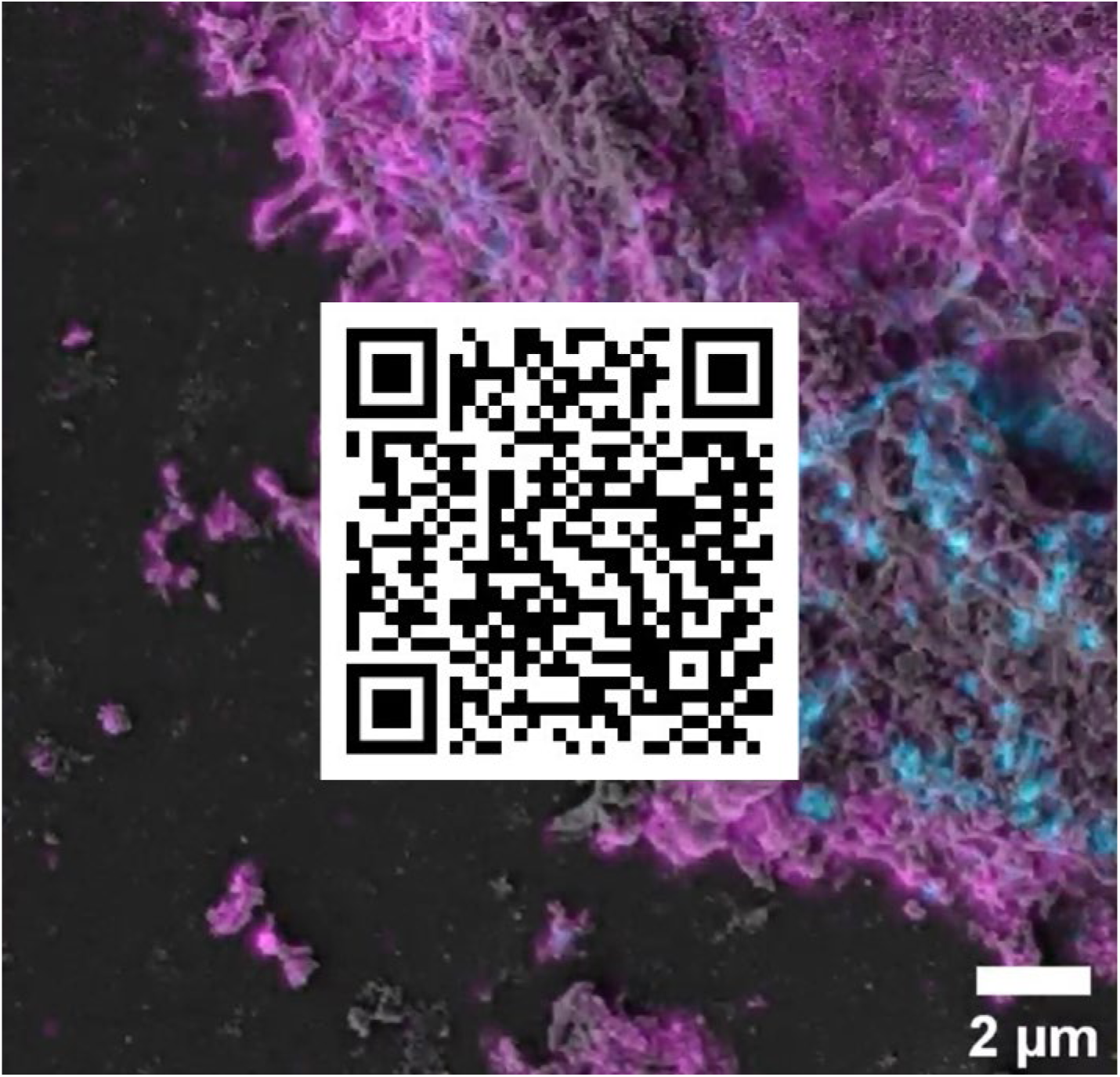
FFPE CLEM imaging over large fields of view.

